# A mechanistic study on the tolerance of PAM distal end mismatch by SpCas9

**DOI:** 10.1101/2023.10.16.562469

**Authors:** Dhritiman Dey, Rudra Chakravarti, Oindrila Bhattacharjee, Satyabrata Majumder, Dwaipayan Chaudhuri, Kazi Tawsif Ahmed, Bireswar Bhattacharya, Anupam Gautam, Rajveer Singh, Rahul Gupta, Velayutham Ravichandiran, Dhrubajyoti Chattopadhyay, Abhrajyoti Ghosh, Kalyan Giri, Syamal Roy, Dipanjan Ghosh

**Affiliations:** National Institute of Pharmaceutical Education and Research-Kolkata; National Institute of Plant Genome Research-Delhi; Department of Life Sciences, Presidency University, Kolkata, India; Department of Botany, Visva-Bharati, Shantiniketan, India; Institute for Bioinformatics and Medical Informatics, University of Tübingen, Sand 14, 72076 Tübingen, Germany; International Max Planck Research School ‘From Molecules to Organisms’, Max Planck Institute for Developmental Biology, Max-Planck-Ring,5, 72076 Tübingen, Germany; Indian Institute of Chemical Biology-Kolkata; Sister Nivedita University-Kolkata; Bose Institute-Kolkata

**Keywords:** CRISPR-Cas9, RMSD, Off-targeteffect, mismatch tolerance, DNA-RNA conformational stability, PAM distal end

## Abstract

CRISPR-Cas9 is the newest technology available for targeted genome editing. It is very efficient and cheap compared to other genome editing techniques. However, its therapeutic application is limited due to its off-target activity. To have a better understanding of this off-target effect, we concentrated our efforts on its mismatch-prone PAM distal end. Current off-target prediction algorithms use RNA-DNA complementation derived energy as a major factor in predicting off-target effect. RNA-DNA complementation derived energy drives Cas9 conformational change, which in turn drives its functional activity. In the case of lower RNA-DNA complementarity, a partial conformational change occurs resulting in a slower reaction rate and partial activity. However, extensive mismatches are often tolerated despite lower complementation derived energy available from RNA: DNA duplex formation. Thus, the off-target activity of Cas9 depends directly on the nature of mismatches which in turn result in deviation of the active site of the enzyme due to structural instability in the duplex strand. In order to test the hypothesis, we have designed an array of mismatched target sites and performed in vitro and cell line-based experiments to assess the effects of PAM distal mismatches in Cas9 activity. For further mechanistic validation, Molecular dynamics simulation was performed and it revealed that certain mismatch mutations induced pronounced conformational instability within the RNA-DNA duplex, leading to elevated root mean square deviation (RMSD) values. We found that, target sites having mismatches in the 18th to 16th position upstream of the PAM showed no to little activity.

## 1. Introduction

CRISPR-Cas9 system is currently the most useful and extensively used system for targeted genome editing. The Cas9 protein from *Streptococcus pyogenes* (SpCas9) binds with guide RNAs (gRNA) to specifically dock to a DNA sequence that is complementary to the probe region of gRNA. The SpCas9, an endonuclease, then cleaves the bound DNA. This simple, easy-to-use system allows for targeting any desired DNA sequence. These DNA target sequences however need to have a Protospacer Adjacent Motif (PAM) which is a tri-nucleotide sequence immediately following the target sequence [1].

In type II CRISPR systems, SpCas9 the multi-domain endonuclease that, along with a single guide RNA (sgRNA), targets specific DNA sequences with a PAM sequence, typically NGG [1]. The sgRNA binds to apo-Cas9, causing a conformational change that activates SpCas9 for DNA binding. The SpCas9-sgRNA complex then stochastically searches and binds to DNA sequences complementary to the sgRNA’s first 17-20 nucleotides at the 5’ end. This binding leads to the cleavage of the target DNA between the 3rd and 4th nucleotide upstream of the PAM, creating a Double-Stranded Break (DSB), which can be repaired via non-homologous end joining (NHEJ) or homology-directed repair (HDR)[2, 3]. The PAM site is crucial for SpCas9 binding initiation, while the seed sequence, located near the 3’ end of the sgRNA, is critical for subsequent SpCas9 binding, R-loop formation, and nuclease activation. The Cas9-sgRNA complex’s crystal structure shows that the HNH and RuvC nuclease domains are responsible for cleaving the complementary and non-complementary DNA strands, respectively [4].

Recent studies have shown that the correct interaction of the gRNA with the targeted DNA sequence is essential and plays a crucial role in the correct functioning of the Cas9 endonuclease activity [5]. Although CRISPR is a fast, efficient, and cheap genome editing technique, its off-target effects, give rise to genetic instability causing unwanted phenotypes thus limiting its application in therapeutic purposes[6, 7]. There are many reasons for the off-target effects of CRISPR systems; the tolerance of mismatches in the PAM distal end being one of the most important one. SpCas9 has been shown to tolerate mismatches in the PAM distal end[8, 9] while being much more conservative in the seed region and its adjacent areas. This tolerance is explained by stabilization of the distorted duplex by a domain loop that penetrates this duplex [10]. Even, the bridge helix arginine-rich domain and REC lobe of the Cas9 protein also holds the key for off-target activities [11].

The aim of this work is to shed light on the effect of the number and position of mismatches in the PAM distal end of target DNA on the functional activity of SpCas9. We have also tried to give a mechanistic insight on the tolerance of various types of mismatches in the PAM distal end on the functional activity of SpCas9 through Molecular Dynamics (MD) simulation. The tolerance profile was found to vary with the nature of mismatch that was attributed towards the stability of DNA-RNA duplex.This variation may also result due to variation in GC content between different sequences[12]. In other words, the significance given to one position may not hold much value when compared across other sequences of varied GC content. Due to this, we have suggested using structural instability of the RNA: DNA duplex in predicting tolerance of mismatches. For this, we have studied the effect on SpCas9 activity upon mismatches and deletion in the complementary PAM distal end of the gRNA probe. Furthermore, we have also studied many varieties of staggered mismatches and their activities using in vitro cleavage assay and cell-based reporter assay. The results from cell-based assays closely mirrored those obtained from our in vitro experiments. We have shown a high tolerance for various mismatches which is difficult to explain based on energy or position. To gain a more comprehensive understanding of our *in vitro* and cell-based findings, we performed molecular dynamics simulations. We have chosen 5Y36 for the study as it is denoted as the cleavage-compatible conformations with the target DNA [13]. Several other structures for the CRISPR-Cas9 system with RNA and DNA are deposited in the PDB but in all of that, residue 840 (mainly His840, which is the residue responsible for the nucleophilic attack to the phosphorus atom by a water molecule) of HNH domain is very distant from the scissile P atom which resides between bases 3 and 4 upstream of PAM. This phosphodiester bond (P-O) is cleaved by one-metal-ion hydrolysis mechanism. 5Y36 is solved with two Alanine substitution mutations in the protein part at position 10 and position 840. Asp10 resides in the RuvC domain which helps in binding two Mg2+ ions along with Glu762, Asp986. Then using two-metal-ion hydrolysis mechanism, the phosphodiester bond between bases −4 and −3 of the non-target strand is hydrolysed by His983 or His982 [14–16]. These two substitution mutations cause loss function in the CRISPR-Cas9. In **Figure1F**, we have depicted the RNA-DNA duplex of the wild type including the cut site phosphate atom. We computed the root mean square deviation (RMSD) of RNA-DNA double helix using all the heavy atoms. We have selected bases 1 to 19 from RNA (chain B) and bases 21 to 39 from DNA (chain C) because this segment comprises the double helix structure. Mutations have been introduced in the region spanning bases 31 to 39 of DNA chain C. Molecular dynamics simulations revealed that certain mismatch mutations induced pronounced conformational instability within the RNA-DNA duplex, leading to elevated root mean square deviation (RMSD) values. Conversely, other mutations had negligible effects on the duplex stability. This contrast in stability directly translated into the observed catalytic activity of the mutants. Specifically, those mutants characterized by a highly stable RNA-DNA duplex exhibited full catalytic activity, while mutants associated with significant duplex instability displayed negligible or no catalytic activity. A subset of mutants demonstrated intermediate stability in the duplex, resulting in partial catalytic activity. This study signifies the correlation between the effects of mismatches, on SpCas9 functional activity with the stability of the DNA:RNA hybrid. This corelation will play a key role for designing sgRNA for high precision genome engineering study and will lead to the development of new sgRNA prediction algorithms.

**Figure 1.**
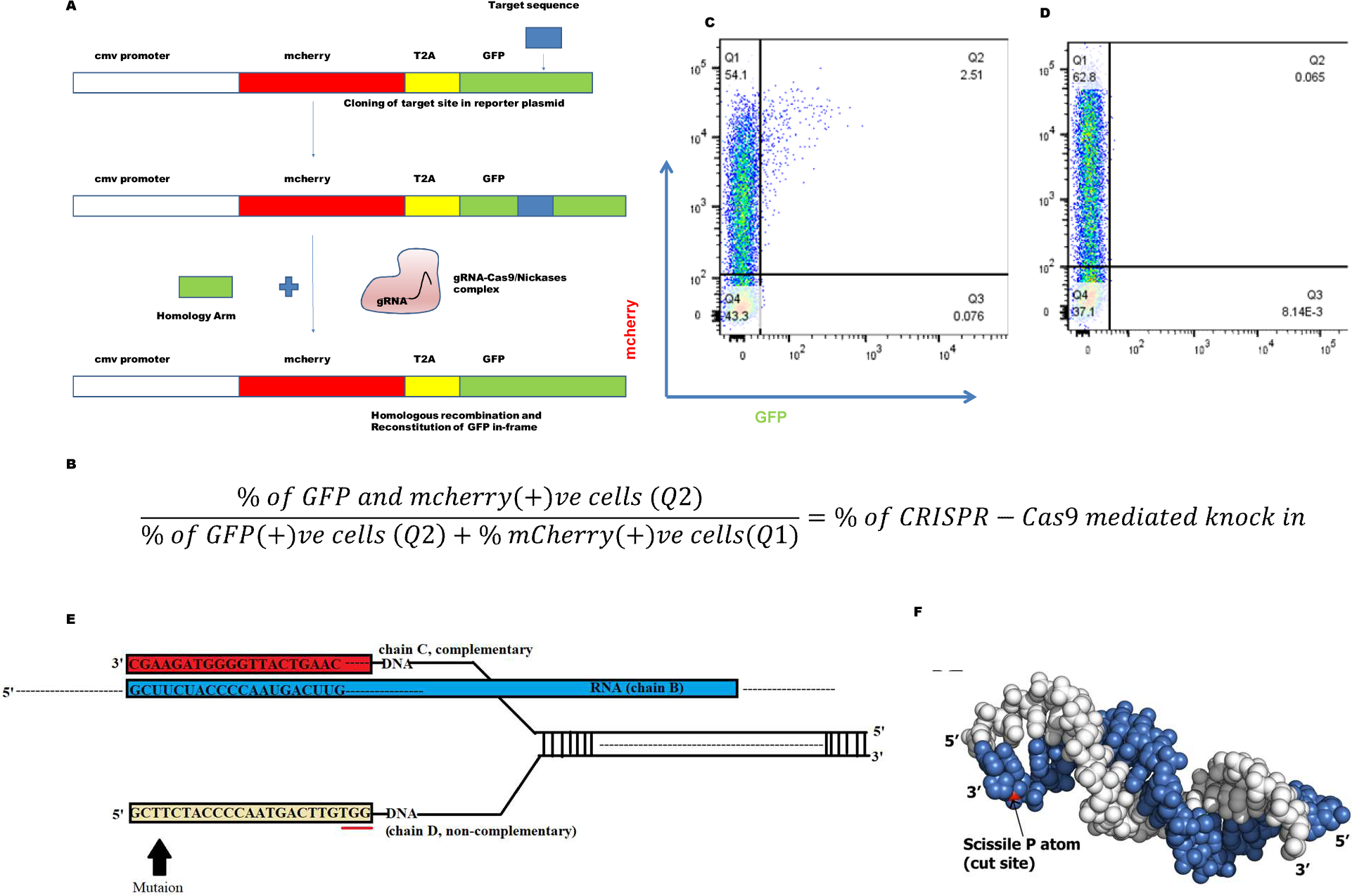
A) Functional module of the reporter assay vector system. B) Equation depicting percentage of knock-in calculated form FACS based experiments C) and D) FACS representation of cell based knock-in. Q1 represents mcherry positive population (transfected cells). Q2 represents dual positive population (HDR knockin population amongst transfected population). Q3 represents GFP positive population and Q4 representes non-transfected cells. E) Sequence composition of the Wild type CRISPR-Cas9 system and site for introduction of mismatch mutation. F) The duplex structure of Wild type, comprising RNA (base 1-19, depicted as white spheres) and DNA (base 21-39, depicted as blue spheres). The phosphorus atom at the cleavage site (scissile P) is shown in red.

## 2. Results

### 2.1. Cas9 activity upon variation in gRNA probe length and GC content of target site-

Our first aim was to study whether energy consideration holds across various sequences for Cas9 activity. We began our investigation by deleting one residue at a time from the PAM distal end and observing its effect on Cas9 functional activity. The target sequence selected had a 50% GC content target site DNA-TS1. Our results demonstrate that SpCas9 activity is sensitive to gRNA probe length modifications in TS1 (**GCUUCUACCCCAAUGACUUG)** from PAM distal end. For getting full activity from SpCas9, up to 18 nucleotide probe length is sufficient. However, there is a severe drop in the activity when the probe length is reduced to 17 nucleotides. Furthermore, when the probe length is reduced to 16 nucleotides or less, the activity is almost abolished (**Figure2a**). However, it is not known whether the gRNA probe is unable to bind the target DNA or post binding doesn’t have sufficient energy to drive the reaction. To ascertain this we performed a pull-down experiment with non-digesting gRNA (16 nucleotides, 15 nucleotides, 2015mm and 2016mm). Our results show that all the gRNAs can bind the target DNA (**Figure2e**). This indicates that there is a cut-off complementarity required. The *in-silico* energy calculation, based on complementarity with DNA target, defines the minimum cut-off energy (ΔG) for this target as -15.6 kcal mol^−1^ (**Table 1**). TS1 with 16 nucleotides has ΔG of-14.2 kcal mol−1, which according to our digestion data is insufficient to drive Cas9 for even minimum activity. However, this value of minimum energy is not applicable for TS2 and TS3, with 80% and 20% GC content respectively (**Table 3**). While in TS3, even after truncating up to 12 nucleotides activity remained. This shows that at much lower energy Cas9 was still active. However, in TS2 this was not the case. We found that Cas9 activity was inconsistent across various truncations of TS2 and TS3 (**Table 2**) (**Figure2c**). This is despite TS2 having 80% GC content. Thus, having sufficient scope for complementation derived energy. It has been reported that if the GC content of the target site is high (80%), then there is an anomalous pattern of Cas9 mediated digestion. The percentage of digestion is reduced drastically. On reducing the gRNA probe length, the GC content of the probe is normalized and we can see incomplete but enhanced digestion (**Figure2d**). This result is at par with the findings of Wang et al. in which they reported about reduced efficiency of Cas9 with gRNA probes having abnormal proportions of GC content [12]. With the above results, we can say that to drive the reaction forward, even to the extent of partial activity, complement length derived energy alone cannot be a sole factor. Even with smaller probe designs, this effect will not be mitigated [17]. We found an erratic pattern with TS4 & TS5, with GC content 40% and 60% respectively, as well (data given in supplementary file S3). Thus off-target predicting algorithms with only energy consideration will not suffice. This simple experiment demonstrates this and clearly shows that selections of target probes, based on energy criteria alone, may be prone to off-target activity. We performed a time point digestion assay to observe the kinetic profile of wild type 20 nucleotides and 17 nucleotides were plotted and examined (**Figure2b**). Results show that the kinetic profile for both cases is very different. In the wild type 20 nucleotide, we can see that the stationary phase is attained very early where a majority of the target DNA is digested. However, in the case of the 17 nucleotides, very little DNA is digested and the rate is slow compared to that of wild type. However, for TS2 and TS3 we find that the digestion is not following the energy derived from complementarity.

**Figure 2.**
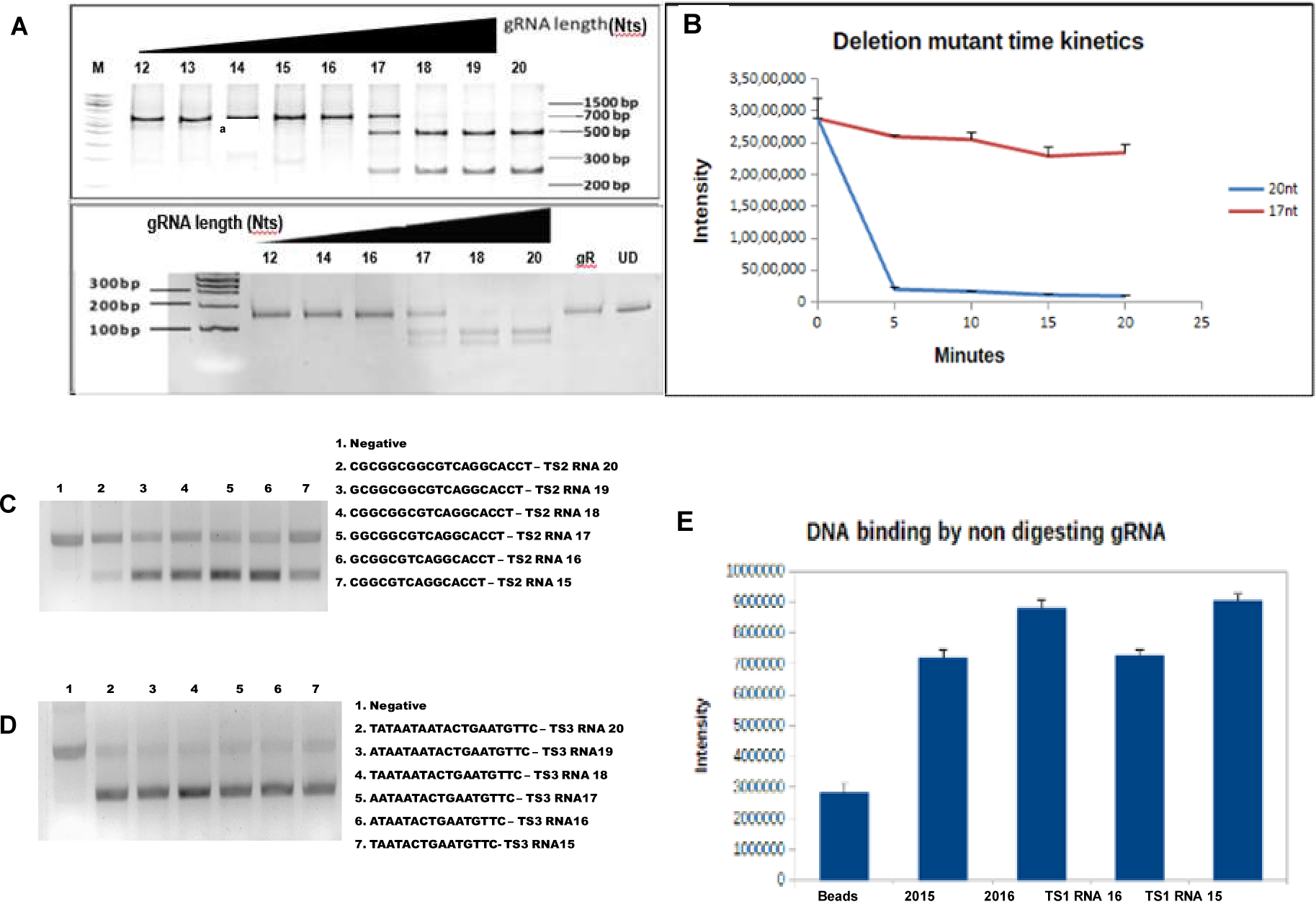
Cas9 functionality upon using truncated gRNA probes. A) DNA digestion with truncated gRNA probe TS1 – DNA digestion is dependent on RNA length. 20nt probe length to 18nt probe length is sufficient to produce digestion. However, at 17nucleotides length, partial digestion is produced. 16 nucleotides or less produced no digestion. B) Timepoint digestion assay of 20 nucleotides and 17 nucleotides. A significant difference between the wild type and truncated gRNA. C) DNA digestion with truncated gRNA probe TS2 – DNA digestion is not dependent on RNA length. D) DNA digestion with truncated gRNA probe TS3 – DNA digestion is again not dependent on RNA length. E) Pull-down of Cas9-gRNA complex using Affinity Chromatography - Pull down of Cas9-gRNA complex with higher bound DNA than control beads indicating the ability of these gRNA to bind their target DNA.

**Table 1.**
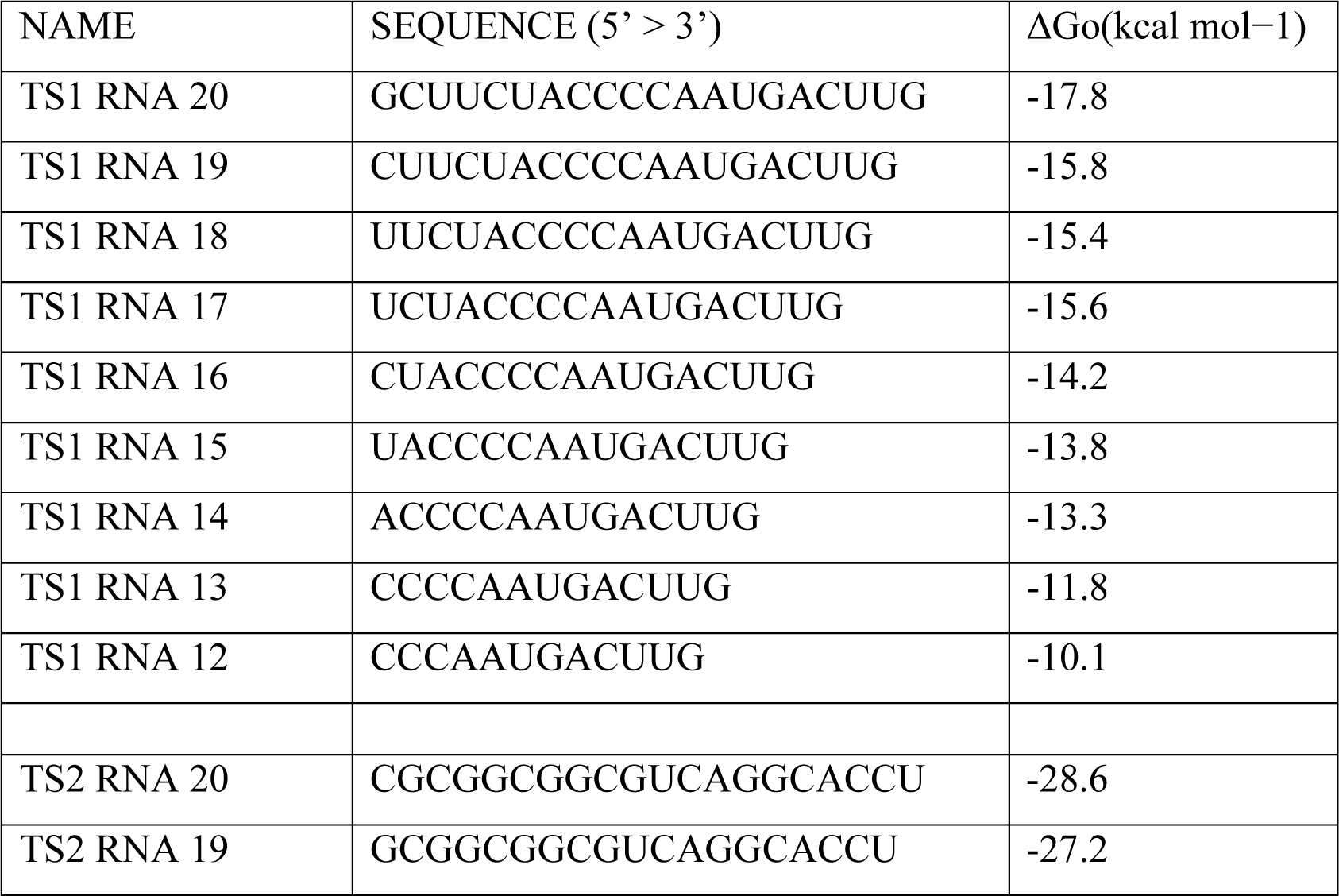
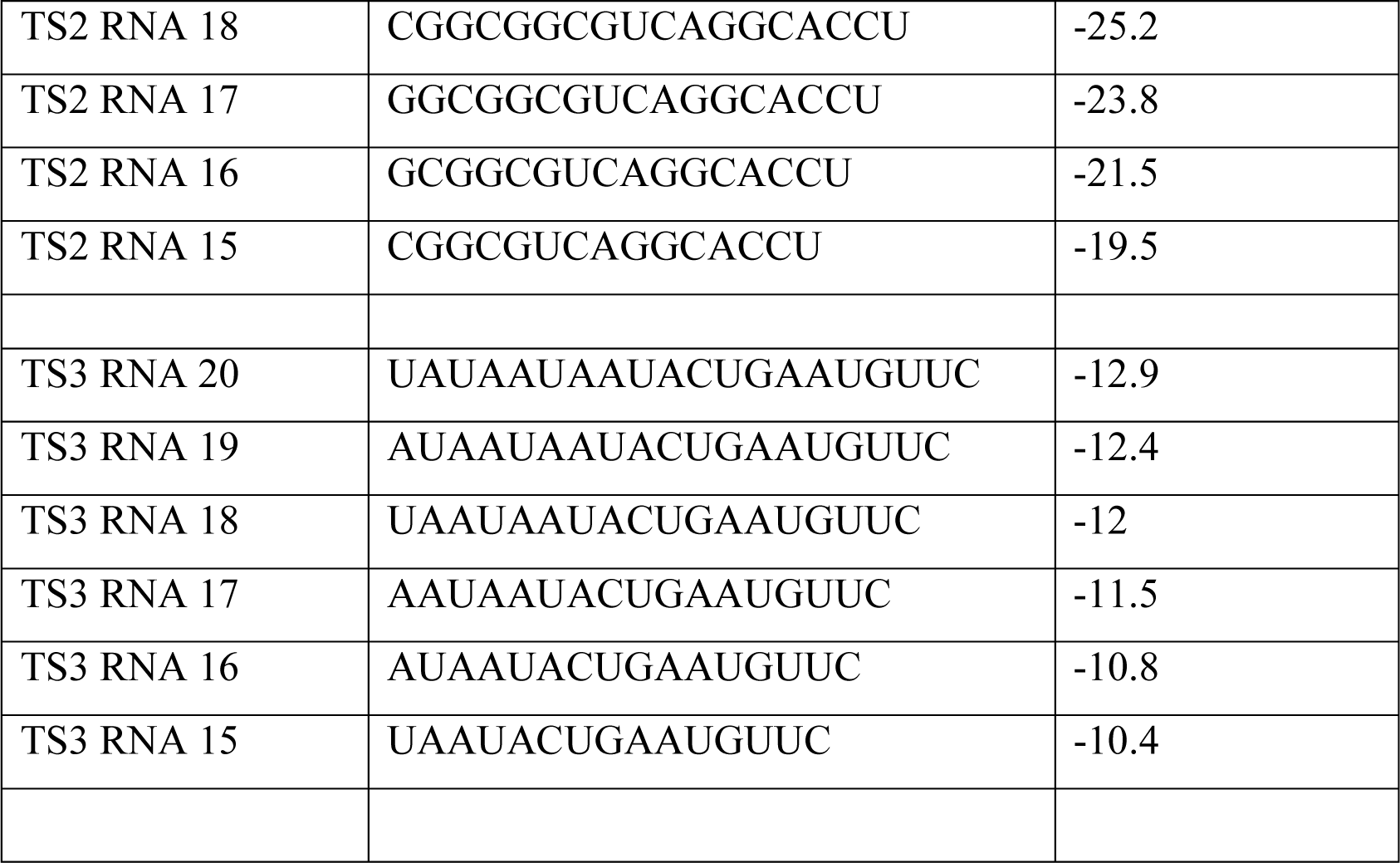
RNA drop.

**Table 2.**
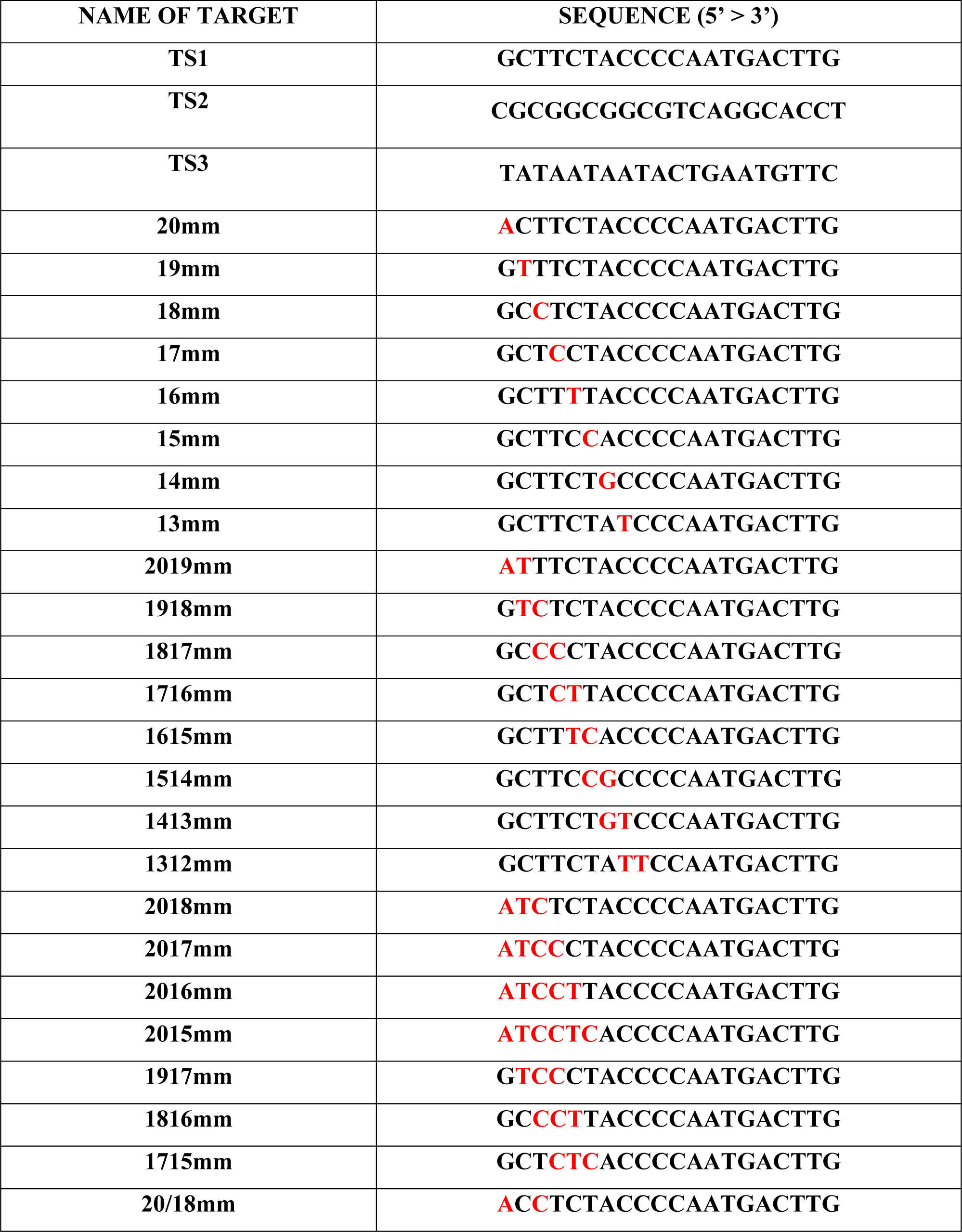
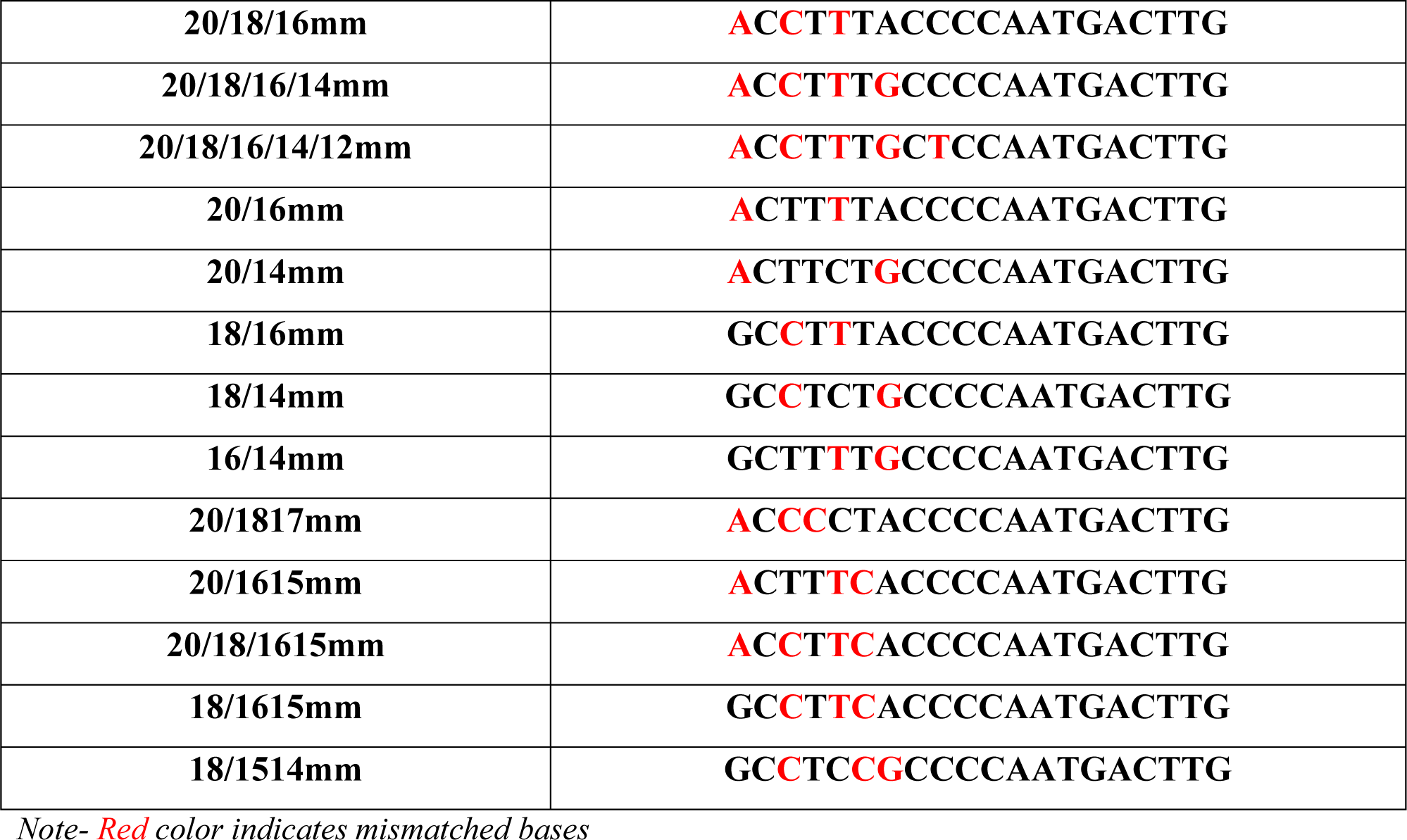
list of Target DNA with mismatches in accordance to TS1.

### 2.2. Single nucleotide mismatches in SpCas9-gRNA complex interaction with targeted DNA TS1

We have observed the effects of nucleotide deletion on the activity of the Cas9:gRNA complex. With different target sites, the behavior of Cas9 was different. It is thus difficult to predict by simply emphasizing the energy derived through complementary binding. Another aspect that has been widely reported as a factor in off-target activities is the position at which the mismatch is occurring. There are reports of the position 17 to 14 being important. However other reports show that the distal end from PAM is mutation prone. Other reports designate 13 or so from PAM distal end as mismatch-prone [18–22].To address this we designed many mismatched target sites derived from TS1. We now kept the length of the probe the same, but sequentially mutated the TS1 site to simulate a variety of mismatches. They were divided into categories, Single mismatch, sequential mismatch, bi sequential mismatch, and staggered mismatches. To study these mismatches – Cas9 digestion assay, Cell based Knock in assay, Cas9 kinetic assay, energy calculations, and finally Molecular dynamics simulation were performed. As per general expectations, single mismatches were easily tolerated (**Figure3a, 3b**). This was reflected in the digestion kinetic profile which showed full activity similar to wild-type (**Figure3c**). The DNA: RNA binding energy was also similar to wild type (**Figure3g**, Supplementary Table S1). Single mismatch mutations did not induce noteworthy structural deviations in the RNA-DNA duplex, as evidenced by the consistently low Root Mean Square Deviation (RMSD) values observed during the entire simulation. The high stability of the duplex conformations allowed the catalytic domain (HNH domain) to execute cleavage without encountering any obstacles, resulting in these mutants displaying full enzymatic activity (> 85% digestion). We conducted structural comparisons between the initial and average conformations (extracted from the 40 ns to 50 ns timeframe) of 18mm (**Figure3e**) and 13mm (**Figure3f**) to visually illustrate the observed deviations.In staggered mismatch mutants, two categories emerged: highly stable duplexes (low RMSD) with high catalytic activity and considerably less stable duplexes (high RMSD) with low catalytic activity. Furthermore, no significant instability was found in the duplex upon single-nucleotide mismatches (**Figure3d**). Our results indicate that single nucleotide mismatches are well tolerated irrespective of their position duplex strand. Near full activity is observed at all the positions in the gRNA probe tail (**Figure3a, b**). This shows that Cas9 can tolerate a single nucleotide mismatch, irrespective of the position of the mismatch in the PAM distal end. If the single mismatch is right at the terminal portion of the PAM distal end then it is best to avoid it by using a truncated gRNA probe. However, if it is further interior then it is best to not proceed as our results indicate there is a possibility of the mismatch being tolerated.

**Figure 3.**
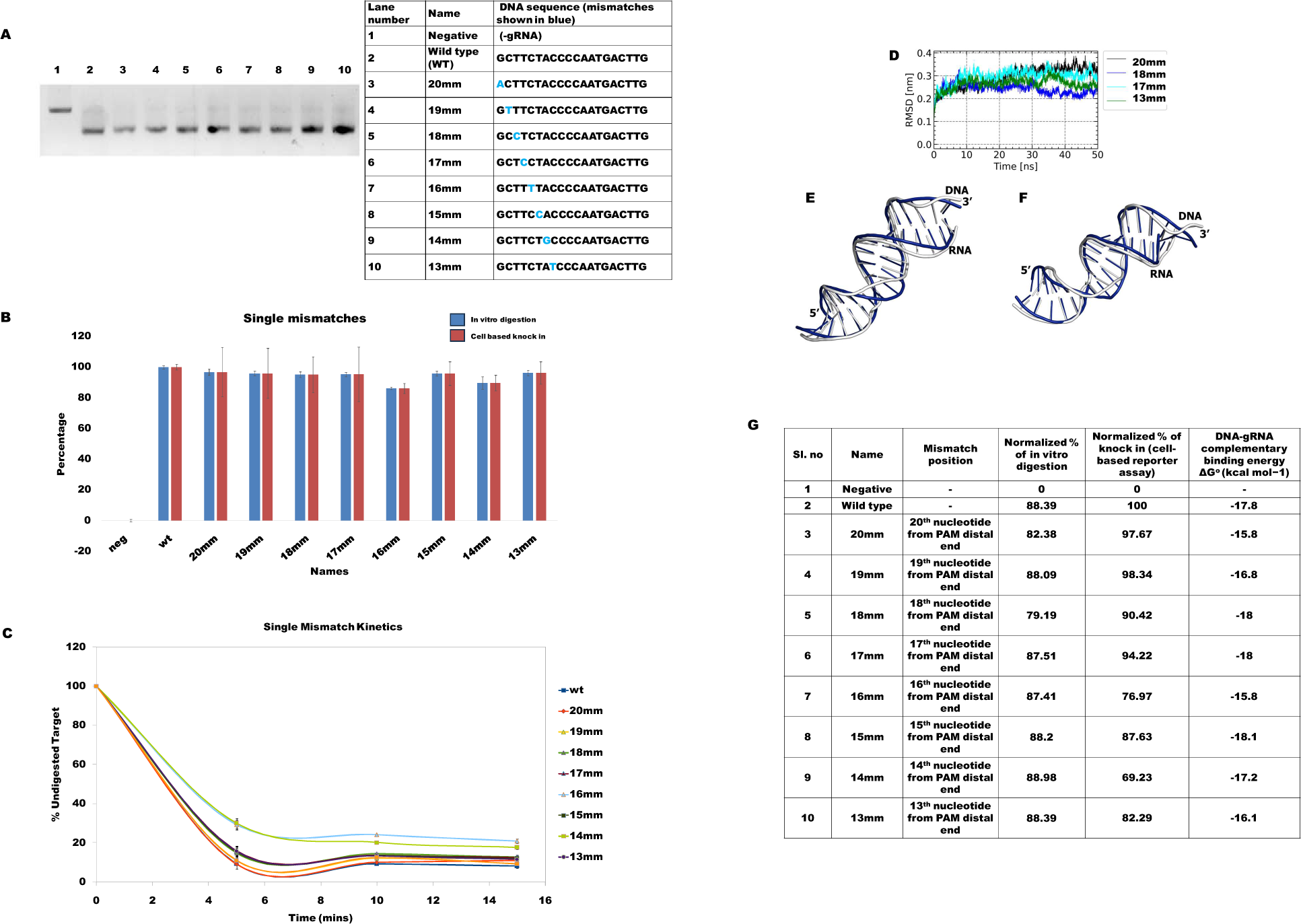
Cas9 functional activity on encountering single mismatches across various positions. A) Cas9 digestion activity of wild type and single positional mismatches in the TS1 target site - Digestion similar to wild type. B) Graphical representation of the correlation between percentage of DNA digestion and percentage of knock-in (cell line based) C) Time-point digestion assay of wild-type and single positional mutants in the TS1 target site - Mutant reaction rate similar to wild-type. D) RMSD trajectory of the RNA-DNA duplex from single mismatch mutants. Superimpostion of initial and average conformation (computed from 40ns-50ns) of RNA-DNA duplex of E) 18mm (RMSD 2.05 Å) and F) 13mm (RMSD 2.4 Å). G) Tabular representation of nature of mismatch, %of undigested target after 15 mins (in vitro digestion), normalized %of knock in (cell based reporter assay), and DNA-gRNA complementation derived energy.

### 2.3. SpCas9-gRNA complex interaction with sequentially mutated nucleotides in TS1

Five Sequential mutations were made from the PAM distal end and three triple sequential mutations were designed for19th position onwards (**Table 1**). This was done to gauge the effect of sequential mutation on Cas9 activity. We found that from sequential tri-nucleotide mismatch to sequential hexanucleotide mismatch there was minimum DNA cleavage activity observed. At 2017mm (17 nucleotide complementarity) partial activity is found while at 2016mm (16 nucleotide complementarity) there is no activity (**Figure4a**). The kinetic profile was also examined and it was observed that the rate was also very different from that of the wild type (**Figure4b**). The reaction profile of sequential mismatch remains stagnant right from the beginning (**Figure4c**). The in-silico energy calculation also shows that the ΔG value for the sequential mismatch is less compared to wild type and perhaps due to this very little digestion occurs (**Figure4g**, Supplementary Table S1). Thus with three or more sequential mutations, the activity of Cas9 is severely affected. Other sequential mutations from the 18th position onwards are also unable to digest. From the 19th position there is a partial digestion. This shows that once triple sequential mutations move away from the Pam distal end they inhibit the activity of Cas9. In the case of sequential mismatches, we observed two distinct scenarios. In one scenario, mutations resulted in significant RMSD values (**Figure4d**), indicating a notable decrease in stability (e.g., 2017mm, 2015mm, 1816mm with mean RMSD values exceeding 3 Å). Consequently, these mutants exhibited no detectable catalytic activity (< 3% digestion). For instance, 2015mm displayed a high mean RMSD value of 3.7 Å and a minimal digestion value of 2.07%. In the other scenario, these mutations did not induce substantial deviations in the duplex structure, leading to partial catalytic activity. For instance, 2018mm had a mean RMSD of 2.8 Å with a digestion value of 55.52%. Upon superimposition, the initial conformation of the RNA-DNA duplex in 2015mm displayed significant structural deviation from the average conformation, with an RMSD of 3.7 Å (**Figure4e**). In contrast, the RMSD for 2018mm was quite low, at 2.5 Å (**Figure3f**) even though the number of mutations in 1816mm and 2018mm is the same

**Figure 4.**
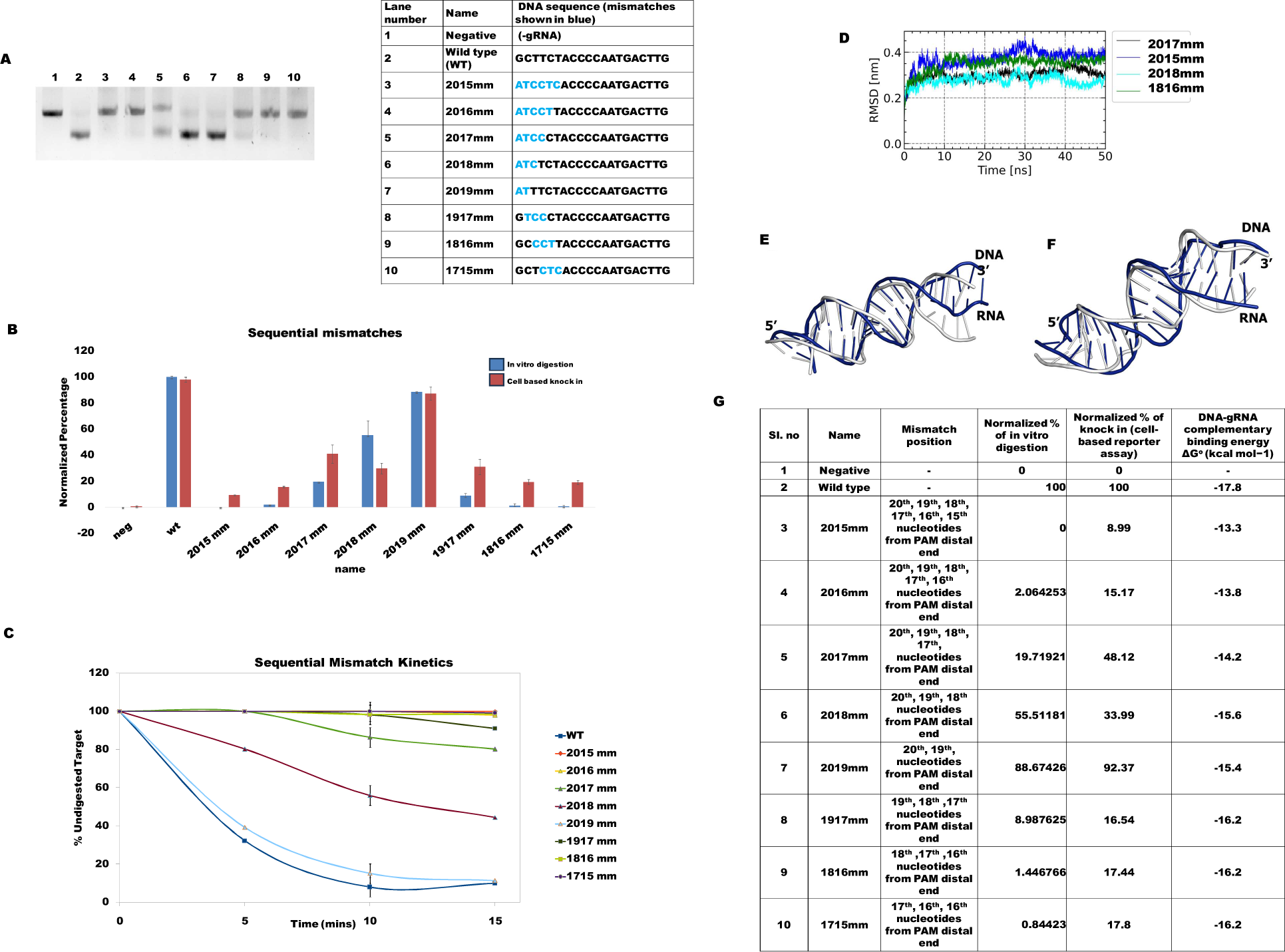
Cas9 functional activity on encountering sequential mismatches across various positions. A) DNA digestion of Cas9 and sequential positional mismatches in the TS1 target site - Cas9 has very little activity when there are multiple sequential mutations. B) Graphical representation of cell based reporter assay system depicting the percentage of knock-in. C) Timepoint digestion assay of Cas9 of wild type and sequential mismatches in the TS1 target site – Significant difference between the wild type and sequential mismatches D) RMSD trajectory of the RNA-DNA duplex from sequential mismatch mutants. Superimpostion of initial and average conformation (computed from 40ns-50ns) of RNA-DNA duplex of E) 2015mm (RMSD 3.2 Å) and F) 2018mm (RMSD 2.5 Å). G) Tabular representation of nature of mismatch, %of undigested target after 15 mins (in vitro digestion), normalized %of knock in (cell based reporter assay), and DNA-gRNA complementation derived energy.

### 2.4. SpCas9-gRNA complex interaction with bi sequentially mutated nucleotides in TS1

The reports of single and sequential mutations are widely reported and are generally expected to be so. Our analysis of the physical effects on duplex strand upon mismatches is also in agreement with the Cas9 activity results. Cas9 behavior upon encountering sequential double nucleotide mismatches has been explored in this study as well. It was shown in the previous section that the mismatch 2019mm located in the region of 20th and 19th nucleotide still retains Cas9 activity. Energy analysis as well as the parameters of physical effect on the DNA strand was also checked. To further this study, we have designed several such mismatches (**Table 3**). Its effect has been studied from the 13th position to the 20th position localized in the PAM distal end. We report that Cas9 activity in positions 19-18 and positions 18-19 is retained. This is not surprising as even the triple sequential mutation from 20-18 base, at the PAM distal end, retained its activity. Physical parameters of the duplex were not significantly altered from the wild-type duplex. However, when we moved further interior of the duplex, the physical parameters of shear, twist, and torsion had significantly deviated values. This showed that due to these mismatches there is a strain on the duplex strand. This is also reflected in the Cas9 activity. Mismatches in positions 17 to 14 are not well tolerated, as reported earlier [17]. Double mismatch in the positions 16th and 15th showed partial activity while the rest showed no activity. Duplex parameters also showed significantly altered values when compared to the wild type. However, 1312mm in position 13th and 12th from PAM distal end showed full activity (**Figure5a, b**). The kinetic profile also showed that the targets having bi-sequential mismatches in the 14^th^ to 17^th^ position remained partially undigested (**Figure5c**). This was unexpected given the fact that this mutation is occurring in the seed region. Given the current algorithm’s practice of assigning importance to mismatch position, this particular mismatch may have been predicted as a non-tolerable mismatch. But as we can see from the Cas9 activity it is well tolerated. Herein lies the importance of studying the stability of the DNA-RNA duplex. Bi sequential mismatch mutants exhibited mean RMSD values ranging from 2.6 Å to 3.6 Å (**Figure5d**) and were classified as displaying partial catalytic activity (digestion values between 40% and 60%). 1615mm and 1312mm (**Figure5e, f**) exhibited good overall structural alignment between their initial and average conformations, with RMSD values of 3.3 Å and 3.0 Å, respectively.

**Table 3.**
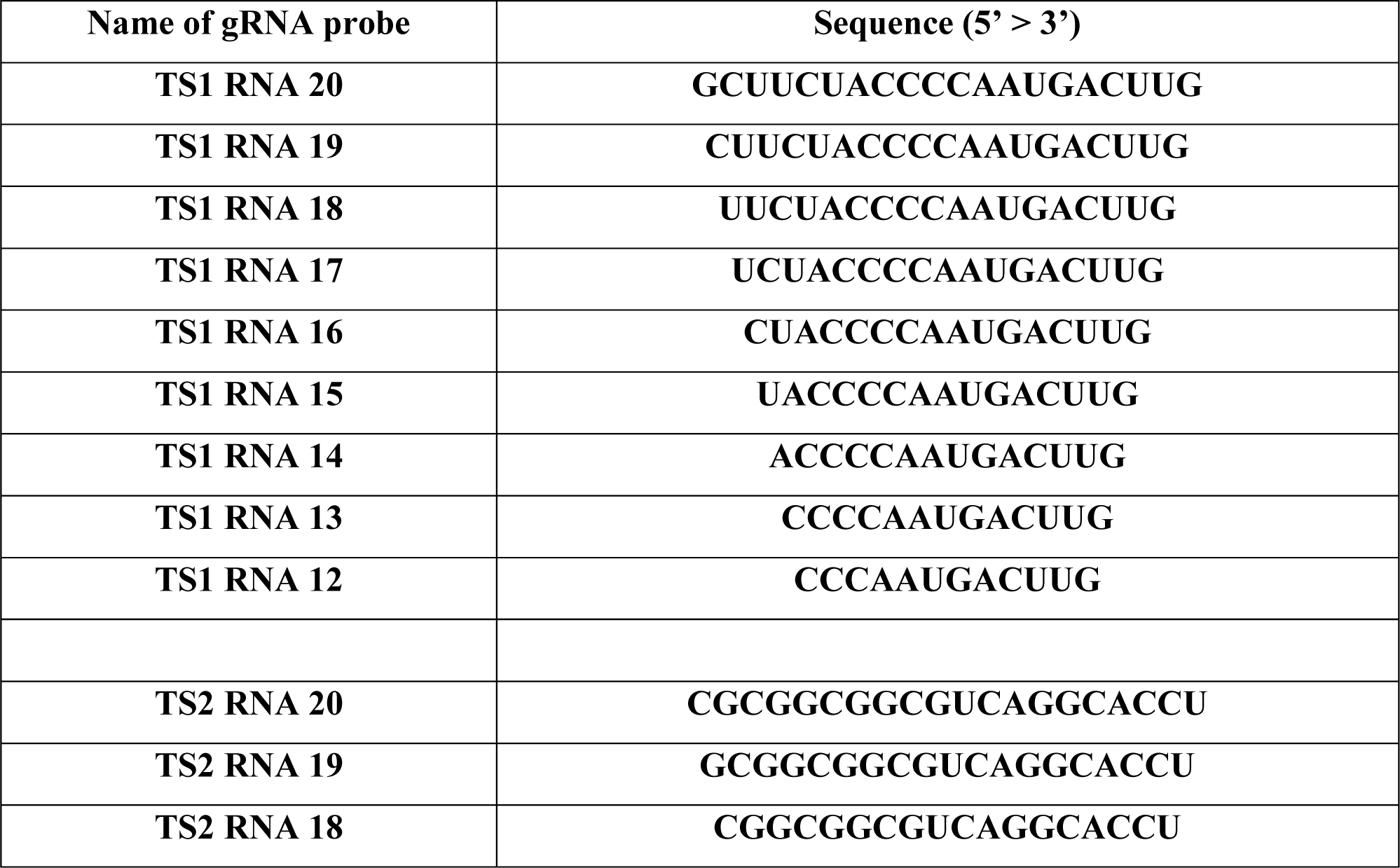
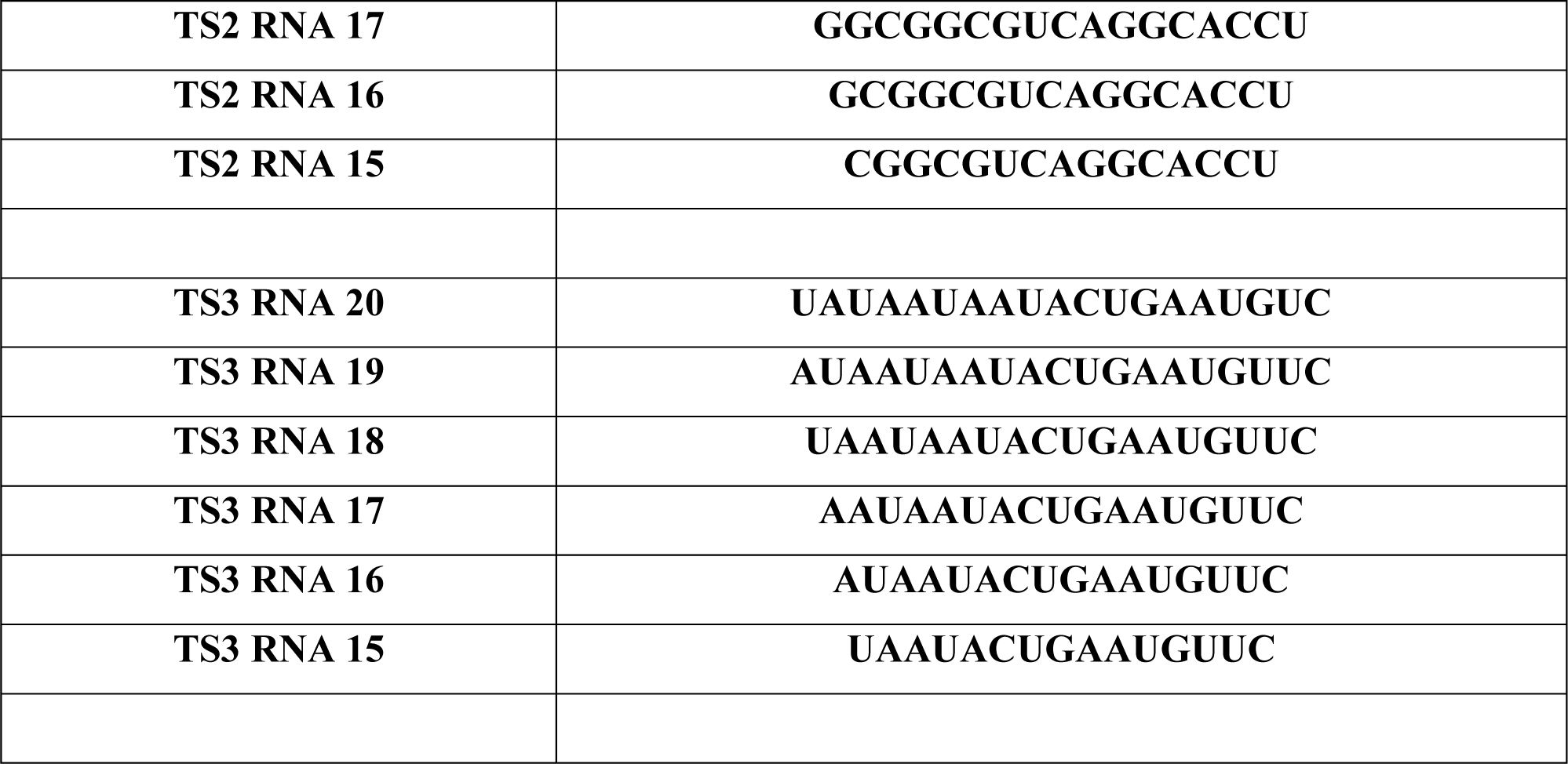
sequences of gRNA probes corresponding to TS1, TS2, TS3, TS4 and TS5.

**Figure 5.**
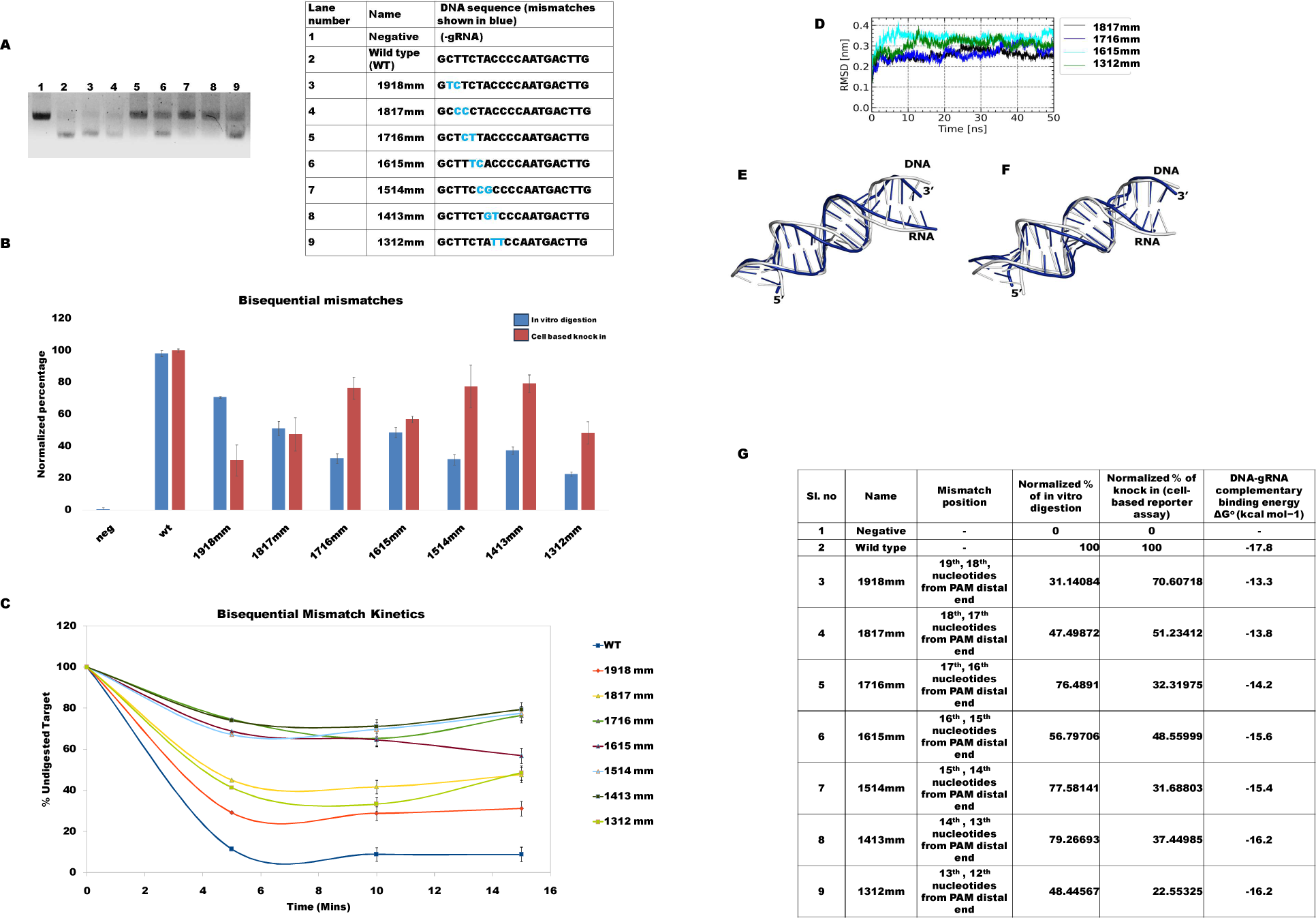
Cas9 functional activity on encountering bi sequential mismatches across various positions. A) DNA digestion of Cas9 with bi-sequential positional mismatches in the TS1 target site B) Graphical representation of cell based reporter assay system depicting the percentage of knock-in. C) Timepoint digestion assay of Cas9 of wild type and bi-sequential mismatches in the TS1 target site D) RMSD trajectory of the RNA-DNA duplex from bisequential mismatch mutants. Superimpostion of initial and average conformation (computed from 40ns-50ns) of RNA-DNA duplex of E) 1615mm (RMSD 3.3 Å) and F) 1312mm (RMSD 3.0 Å) G) Tabular representation of nature of mismatch, %of undigested target after 15 mins (in vitro digestion), normalized %of knock in (cell based reporter assay), local base pair parameters and DNA-gRNA complementation derived energy.

### SpCas9-gRNA complex interaction with staggered mutated nucleotides in TS1

The above mismatches reported in this study have been shown by other researchers as well [8, 9]. we designed staggered mismatches i.e. mismatches that are separated from each other by various distances (**Table 3**). There are not many studies regarding these mismatches and they will serve as rigorous testing for the utility of duplex physical parameters. The various mismatches have been listed in the table. Mismatches 20/18mm, 20/16mm and 20/14mm show Cas9 functional activity. It is interesting to note that the mismatches are getting tolerated even with increasing encroachment to the interior. Mismatches 18/16mm, 18/14mm are partially tolerated by Cas9, however 16/14mm is not tolerated by Cas9 (**Figure6 a, b**). This is hard to explain in terms of assigned positional significance. For instance, 20/18mm exhibited a mean RMSD value of 2.4 Å and a digestion activity of 91%, whereas 18/16mm displayed a mean RMSD value of nearly 4 Å with 0% digestion activity (**Figure 6d**). The structural alignment of these two mutants also supported these findings (**Figure6e, f**).

**Figure 6.**
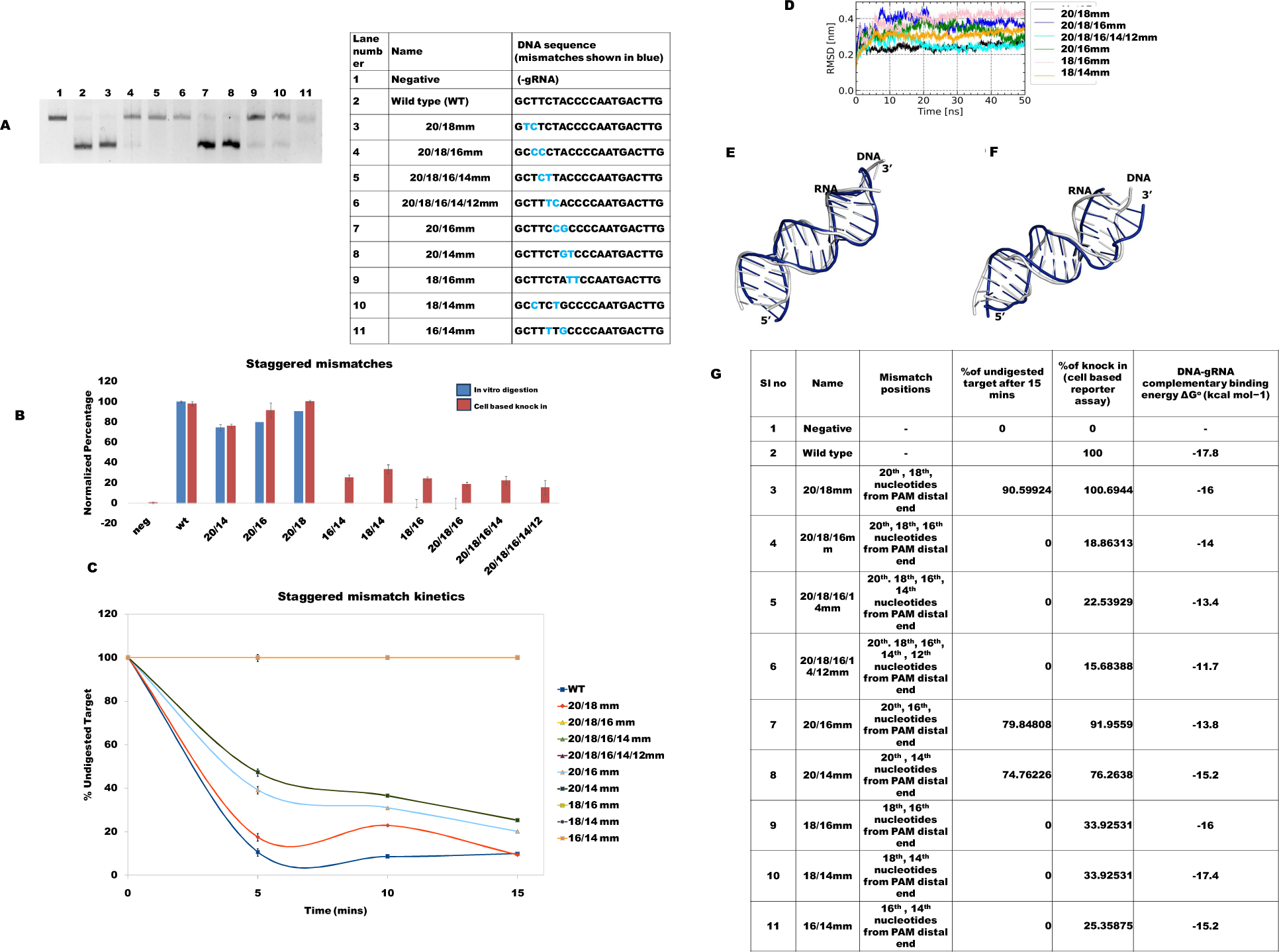
Cas9 functional activity on encountering staggered mismatches across various positions. A) DNA digestion of Cas9 with staggered positional mismatches in the TS1 target site B) Graphical representation of cell based reporter assay system depicting the percentage of knock-in. C) Timepoint digestion assay of Cas9 of wild type and staggered mismatches in the TS1 target site (Since 20/18/16 mm, 20/18/16/14 mm, 20/18/16/14/12 mm, 18/16 mm, 18/14mm and 16/14 mm remained practically undigested throughout the time points they overlapped in the figure) D) RMSD trajectory of the RNA-DNA duplex from staggered mismatch mutants. Superimpostion of initial and average conformation (computed from 40ns-50ns) of RNA-DNA duplex of E) 20/18mm (RMSD 2.43 Å) and F) 18/16mm (RMSD 4.06 Å). G) Tabular representation of nature of mismatch, %of undigested target after 15 mins (in vitro digestion), normalized %of knock in (cell based reporter assay), local base pair parameters and DNA-gRNA complementation derived energy.

We now wanted to see the effect when single and bi-sequential mismatches occur in the same target site. Mismatches 20/1817mm, 20/1716mm, and 20/18/1615mm were studied for their Cas9 activity. We found that all the mismatched sites had activity. This is unexpected as it is in the nucleic acid binding domains region [17]. 20/1817mm, and 20/1716mm gave partial activity, while 20/18/1615mm gave almost full activity (**Figure7 a, b, c**). This was very interesting, especially the tolerance of 20/18/1615mm. Energy and previous positional reports are unable to explain this satisfactorily (**Figure 7g**). Staggered gap mismatch mutants (20/1817mm, 20/1716mm, 20/18/1615mm) fell into categories of partial catalytic activity, with digestion values ranging from 0% to 50%. The mean RMSD values for these mutants ranged from 3.0 Å to 3.6 Å, indicating low to moderate stability (**Figure7d, e, f**).

**Figure 7.**
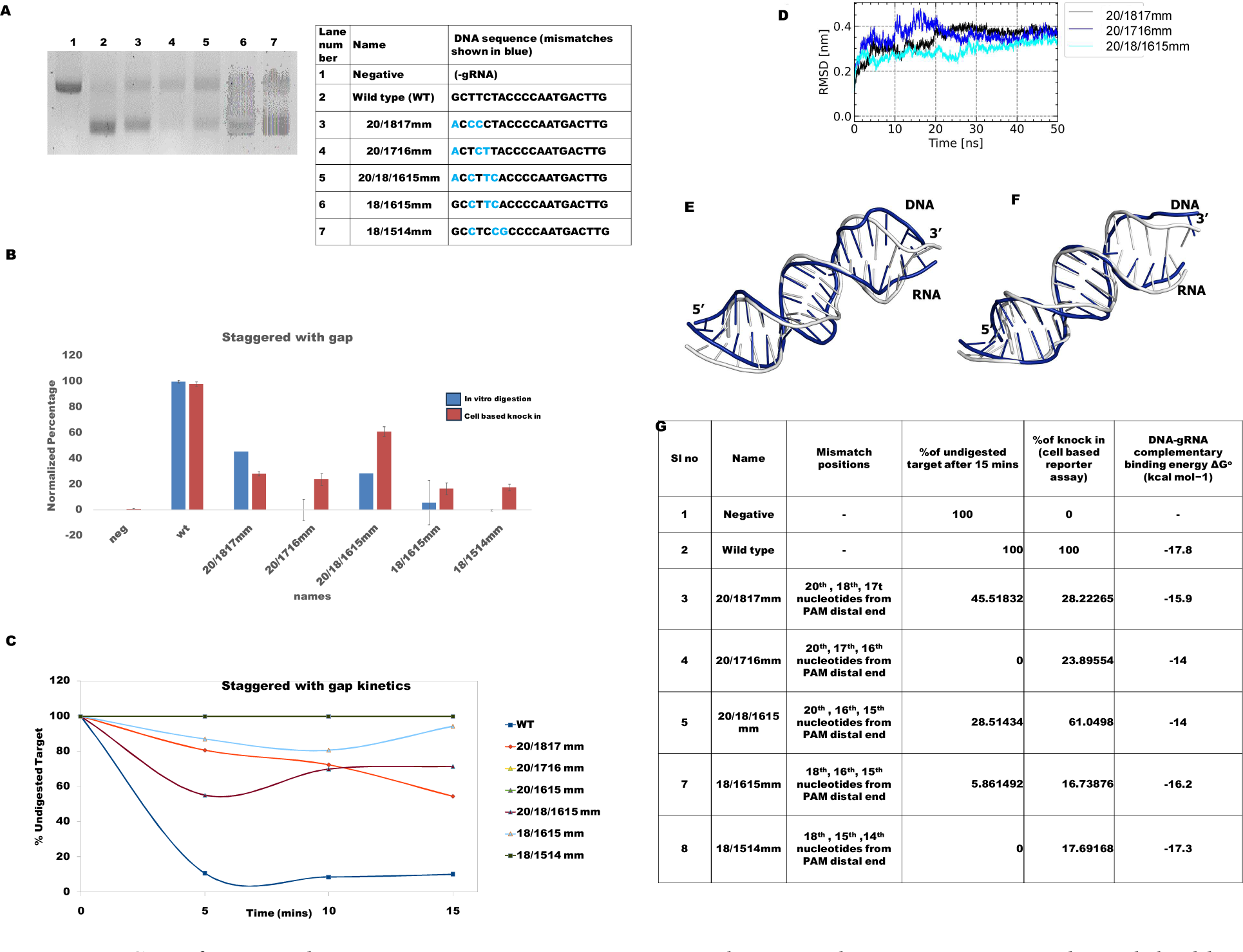
Cas9 functional activity on encountering staggered mismatches containing single and double mismatches across various positions. A) DNA digestion of Cas9 with staggered positional mismatches containing single and double mismatches across various positions in the TS1 target site B) Graphical representation of cell based reporter assay system depicting the percentage of knock-in. C) Timepoint digestion assay of Cas9 of wild type and staggered mismatches containing single and double mismatches in the TS1 target site (Since 20/1716mm, and 18/1514 mm remained practically undigested throughout the time points they overlapped in the figure) D) RMSD trajectory of the RNA-DNA duplex from staggered gap mismatch mutants Superimpostion of initial and average conformation (computed from 40ns-50ns) of RNA-DNA duplex of E) 20/1716mm (RMSD 3.5 Å) and F)20/18/1615mm (RMSD 3.1 Å). G) Tabular representation of nature of mismatch, %of undigested target after 15 mins (in vitro digestion), normalized %of knock in (cell based reporter assay), local base pair parameters and DNA-gRNA complementation derived energy.

## 3. Discussion

CRISPR-Cas9 has shown immense potential in targeted genome editing but it is still a long way from being applied therapeutically. While the generation of “knock-out cells” using CRISPR-Cas9 technology has become routine practice in modern cellular biology, it is mainly restricted to *in vitro*models. Applications of CRISPR-Cas9 have been done in the mice model; however, its efficacy is low and is prone to complications[23, 24]. Non-specific cleavage of genomic DNA can cause cellular toxicity [25]. Their subsequent repair often results in deletions, inversions, and translocations [26–29]. These mutations may cause modulation of gene expression of non-targeted essential genes in the human cell.

Many attempts have been made to modify the Cas9-gRNA complex to reduce the undesired off-target activity. However, such modifications have a sequence-dependent effect on SpCas9 activity [12, 18, 30–32]. Moreover, the total GC content of the target DNA-gRNA hybrid had significant effect on the SpCas9 activity. It was found from our experiments that for target DNAs having abnormally high or low GC content (<20% and >80%) reducing the length of the gRNA probe even below 17 nucleotides retained activity. In modified Cas9-gRNA complexes the off-target effect varies with the target DNA sequences but uniform effectiveness is desired for therapeutic application. For this, we need to be able to predict probable off-target possibilities reliably. Mismatches at the seed region of the Target DNA are not tolerated, whereas from our data it was found that mutations up to certain limits are tolerated at the PAM distal end and thus have a significant implication on the off-target effect of SpCas9. There are many recent off-target probability prediction algorithms including CCTop, CFD, CROP-IT, DeepCRISPR, Elevation, CRISPR-OFF, CRISPR, iGWOS, and uCRISPR[33–40]. Out of these DeepCRISPR, iGWOS and Elevation are based on machine learning, while the rest are based on energetic analysis and experimental rule-driven. However, none of these algorithms take into account the role of DNA-gRNA structural stability that arises due to mismatched target site. In this present work, we attempted to find the role of DNA: RNA complementation derived energy and structural stability on their functional activity. We employed various PAM distal mutations in the target DNA and evaluated the roles of these mismatches on SpCas9 functionality using in vitro cleavage assay as well as cell-based reporter assay. We got very good correlations among them. In vitro assays have the benefit of simplification of complex phenomena and allow to avoid confounding factors, but they do not necessarily reflect the actual processes operating inside a cell. However, the cell-based experiment data also supported and endorsed in vitro data.

Fu, Y., et al., have shown that shorter probe length leads to less off-target effect as less energy is available. They have shown that 16 nucleotides result in no activity in concurrence with our result. A single base loss (via mismatch or deletion) is poorly tolerated when the probe length is 17 nucleotides [17, 41]. On the other hand, when the probe length was 20, single nucleotide mismatch tolerance occurred in the tail region across all positions. However, when the GC content was altered Cas9 activity altered despite shortening the length of the gRNA probe, by previously reported results [12]. Single nucleotide mismatch has also been previously reported to be better tolerated [8, 9, 42]. Many CRISPR-Cas9 off-target evaluators additionally focused on the position of the mismatch and closer the mismatch is towards the PAM proximal end, lesser the probability of off-target effect [43–51]. We observed single mismatches in the PAM distal end and they do not have any significant difference in activity among them. Thus, there is no positional significance or weight attached to single mismatches occurring in this region. We also observed that bi-sequential mismatches were tolerated when they were present in the end positions but when the same was poorly tolerated when the mismatches were present in the 18-16th position from the PAM distal end. However, multiple sequential mismatches are poorly tolerated and upon an increase in mismatch to more than three nucleotides, there is a complete loss of activity which is at par with the previous reports by Ricci, C.G et. al and Hsu, P et al [9, 52]. It was observed from the data that 3 consecutive mismatches is poorly tolerated when the position of the mismatch is in the region of 18-16 nucleotides upstream of the PAM, however similar number of mismatches in the extreme end i.e. in the region of 20-18th position upstream of the PAM was tolerated and Cas9 remained functional. To highlight the importance of DNA-RNA duplex stability or instability in Cas9 activity, we designed a wide variety of mismatches that were staggered (gaps between mismatches). We found that some of these mismatches to be tolerated by Cas9 which could not be explained based on energy or positional effect. We have used MD simulation to characterize how specific mutations in the DNA sequence impact the function of CRISPR-Cas9. The choice of the 5Y36 structure as a model was significant because of its cleavage compatible conformation. One of the key takeaways was the effect of these mutations on catalytic activity. We have computed the RMSD of RNA-DNA duplex from the trajectory as a measure of duplex stability. When single mismatch mutations were introduced, they didn’t disrupt the structure of the RNA-DNA duplex significantly as indicated by low RMSD values. This was at par with our pulldown assay findings. This meant that the system remained stable, and the CRISPR-Cas9 enzyme could carry out its cleaving function efficiently, with digestion rates exceeding 85%. In essence, minor errors in the DNA sequence were tolerated quite well. However, things got more interesting when sequential mismatch mutations were examined. In some cases, these mutations led to noticeable structural deviations in the duplex, reducing stability. As a result, the mutants displayed little to no catalytic activity. For instance, 2015mm had a high RMSD value and a minimal digestion rate. The staggered mismatch mutants revealed another intriguing aspect. They fell into two categories: highly stable duplexes with high catalytic activity and considerably less stable duplexes with low activity. 20/18mm, for instance, had a low RMSD and a high digestion rate, while 18/16mm had a high RMSD and no digestion activity. Staggered gap mismatch mutants also showed varying catalytic activities, ranging from no activity to partial activity. The RMSD values suggested moderate stability in these cases. We established a linear correlation between the average RMSD values and the digestion percentage (catalytic activity). The calculated Pearson’s correlation coefficient (r) is -0.57 (**Figure8**). The computed r value is significant in both p<0.05 and p<0.01. This finding suggests that an elevated RMSD value signifies greater inherent conformational instability within the RNA-DNA duplex structure, leading to a reduction in catalytic activity. Conversely, a lower RMSD value is associated with stable conformation and higher catalytic activity. Higher stability leads to the precise positioning of the scissile P atom in the catalytic site of HNH domain and vice versa.

**Figure 8.**
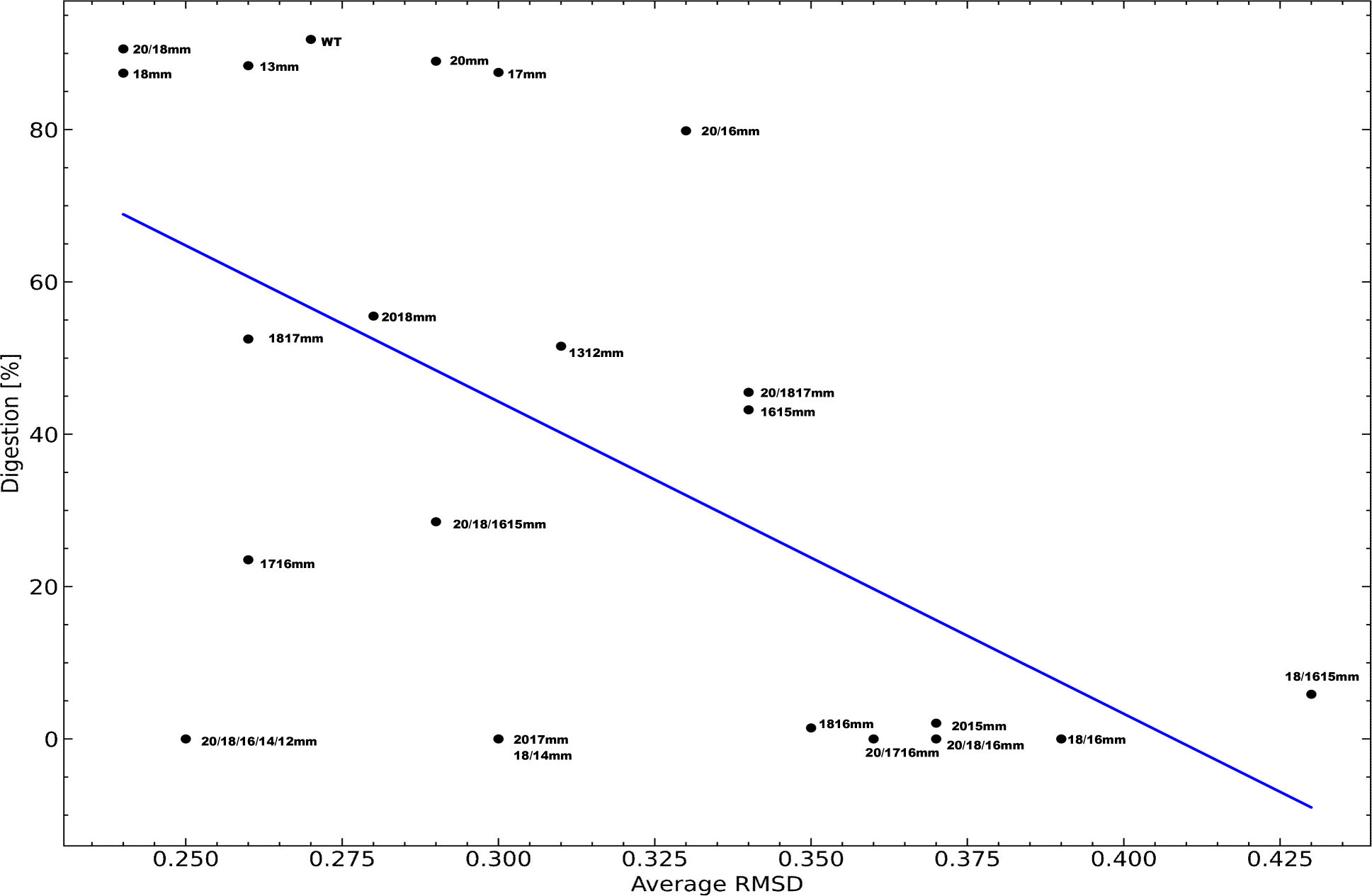
A linear correlation plot that showcases the connection between the mean Root Mean Square Deviations (RMSDs) of the mutant DNA-RNA duplex and the corresponding digestion percentages (catalytic activity).

Overall, a significant correlation was established between the RMSD values and catalytic activity. This indicates that higher RMSD values, reflecting structural instability, were associated with reduced catalytic activity, while lower RMSD values indicated stability and higher activity. Importantly, the study also assessed the conformational dynamics of the RuvC and HNH domains. It found minimal conformational deviations, suggesting that the mutations in the DNA sequence did not significantly alter the dynamics of these catalytic domains. We observed that staggered mismatches having 3 mismatches but with at least 2 nucleotide gaps between the mismatches were tolerated pointing to the fact that the effect of mismatches is limited to the mutated base only. Our work showed that along with the energetics requirement and mismatch position, DNA-RNA duplex instability arising out of mismatches should also be taken into account for reducing the off-target SpCas9 activity.

## 4. Material and Method

### 4.1. Cloning of Target sites

Target sites (containing various mismatches-**Table 2**) primers and their complimentary oligonucleotides were annealed by heating them at 95oC and gradually reducing the temperature by 4oC in a touchdown manner where each cycle is of 4 minutes. The final temperature is 4oC at which the annealed product is obtained and phosphorylated. ESSA plasmid was sequentially double digested with EcoRI and Xho1 at 370C for 2 hours and dephosphorylated. The digested gel product was purified and ligated with 1:100 dilution of the annealed product using T4 DNA ligase. The ligation reaction was performed overnight at 16°C. The ligated product was then transformed into XL1B cells with ampicillin selection. The obtained colonies were screened by sequencing the plasmids isolated from them. After successful cloning of the target sites in ESSA plasmid the target DNA for in vitro experiments were obtained using PCR amplification with HA200F and HA200R primers.

### 4.2. Cloning of probe region in gRNA

Probe regions and their complementary primers (**Table 3**) were annealed by heating them at 95oC and gradually reducing the temperature by 4oC in a touchdown manner where each cycle is of 4 minutes. The final temperature is 4oC at which the annealed product is obtained and phosphorylated. pT7gRNA plasmid was digested with BsmBI at 55oC for 2 hours and dephosphorylated. The digested gel product was purified and ligated with 1:100 dilution of the annealed product using T4 DNA ligase. The ligation reaction was performed overnight at 16oC. The ligated product was then transformed into XL1B cells with ampicillin selection. The obtained colonies were screened by sequencing the plasmids isolated from them.

### 4.3. Synthesis and purification of gRNAs

The cloned plasmids were digested with BamHI and the digested product was purified by gel purification using a Qiagen kit. The linearized vector was then used in an IVT reaction using T7 RNA polymerase to produce gRNA. The produced gRNA was purified using a Qiagen RNA purification kit after DNase I digestion. The purified gRNA was checked for integrity on an 8% Urea Page.

### 4.4. Expression of SpCas9

pet28b Cas9 His was obtained from add gene (Plasmid no. 47327) and was transformed in BL21DE3 Rossetta please cell with kanamycin resistance. The transformed cells were inoculated in LB overnight at 37oC. The overnight cells were again inoculated in a 1-liter culture. Upon reaching a growth O.D. of 0.6, the cells were induced with IPTG at a final concentration of 1 mM. The induced cells were grown overnight at 17oC and after checking for induction the cells were pelleted and proceeded for the purification step.

### 4.5. Purification of SpCas9

The pelleted induced cells were resuspended with lysis buffer (50m Tris pH 8.0, 150 mM NaCl, 10 mM MgCl2, 0.1% NP 40) and incubated in ice for 30 mins. Post-incubation the cells were sonicated and centrifuged at 16000g for 30 minutes at 4oC. The supernatant was collected and charged on a Ni-NTA column at 4oC. The column was washed with various concentrations of Imidazole. The proteins were then eluted at 300mM imidazole and the eluted protein was dialyzed in dialysis buffer (50m Tris pH 8.0, 150 mM NaCl, 10 mM MgCl2, 0.1% NP 40, and 20% glycerol). Dialysis was performed at 4oC overnight.

### 4.6. SpCas9 functional assay

We first synthesized different-sized gRNA by in vitro transcription. The RNA was then purified and checked for integrity on a urea-PAGE. The three individual components i.e. the SpCas9 protein, gRNA, and target DNA were reconstituted and incubated for 60 minutes at 37oC. After incubation was done the reaction was stopped by incubating the mixture at 65oC for 15 minutes. Post this, the samples were first incubated with RNase A at 37oC for 15 minutes. After this, they were incubated with Proteinase K at 55oC for another 15 minutes. Finally, the samples were then run on a Native PAGE gel (for 706 base pair target – 8% PAGE and 122 base pair target – 20% PAGE).

### 4.7. SpCas9 time-dependent activity assay -

The three individual components i.e. the SpCas9 protein, gRNA, and target DNA were reconstituted and incubated for various time points at 37oC. After incubation was done the reaction was stopped by incubating the mixture at 65oC for 15 minutes. Post this the samples were first incubated with RNase A at 37oC for 15 minutes. After this, they were incubated with Proteinase K at 55oC for another 15 minutes. Finally, the samples were then run on a 1% Agarose gel. Their intensity was measured through the image quantification software of Image lab (version 6.0.1)Biorad. The reaction was performed in duplicate and the percentage undigested target DNA was plotted in a graph.

### 4.8. Designing of Cell based reporter assay system

A cell based reporter assay plasmid was constructed such that m-cherry and the first 474 nucleotides of GFP are presented in frame under CMV promoter. Then it is followed by a stop codon and the rest of the nucleotide sequence of the GFP is present. EcoRI and XhoI digestion sites are present flanking the stop codon and between these two restriction sites, the desired target sites are cloned. The reporter assay plasmid is designed such that mCherry is always expressed upon transfection but GFP is not expressed as such. Upon co-transfection of the homologous sequence which is a portion of the in-frame GFP spanning the target and CRISPR system (Cas9 and corresponding gRNA to the specific targets) the entire sequence of GFP comes in frame (homologous recombination) and only then GFP is expressed. Thus, cells which express both mCherry and GFP (dual expression) serve as positive control which indicate successful CRISPR-Cas9 mediated homologous knock-in (**Figure1a**).

### 4.9. Transfection and Flowcytometry analysis-

HEK 293 cells were maintained in DMEM High Glucose supplemented with 10% FBS and 1 % Antibiotic-Antimycotics. The aforementioned reporter assay plasmid containing the various target sites, Homologous arm donor construct and plasmid encoding gRNA and Cas9 were co-transfected using Lipofectamine 2000 (Invitrogen). The amount of the various DNAs was standardized previously and was added in corresponding wells in a 24 well tissue culture plate. Following the addition of the DNAs, the lipofectamine solution (1:100) in opti-mem media (Gibco) was added to each well and at last, 105 HEK 293 cells were added to each well. 4 hours post transfection, the entire opti-mem media was discarded and fresh DMEM High glucose media was added. The transfected cells were maintained for 48 hours and then Flowcytometric analysis of homologous Knock-in was performed using FACS (LSR Fortessa, BD bioscience). The cells that were transfected with the reporter assays system expressed mcherry that was confirmed by the population observed in quadrant Q1 (**Figure1c**). The sub-population of transfected cells in which homologous recombination occurred due to SpCas9 mediated DNA double strand break and repair expressed GFP. this was evident from the dots observed in quadrant Q2 (**Figure1d**). The frequency of Knock-in was calculated using the formula mentioned in **Figure1b**.

### 4.10. DNA pull-down with SpCas9-gRNA complex -

The three individual components i.e. the SpCas9 protein, gRNA, and target DNA were reconstituted and incubated for 30 mins. After that Ni-NTA beads, equilibrated in reaction buffer, was added in equal amount to all the reaction. This was further incubated for another 30 mins at room temperature. Post incubation, the beads were washed twice with reaction buffer. 15ul of reaction buffer was added to all the beads and heated to 65oC for 15 minutes. The samples were first incubated with RNase A at 37oC for 15 minutes. After this, they were incubated with Proteinase K at 55°C for another 15 minutes. The samples were then run on an 8% native page and their intensity was plotted. For the bead control set, gRNA without probe was used. The reactions were done in duplicate.

### 4.11. In-silico energy analysis -

We have calculated delta G (ΔG°) or Gibbs free energy for the guide RNA (gRNA) by utilizing the nearest-neighbor (NN) model which predicts DNA/RNA thermodynamics by utilizing energy values for different base pair motifs. We have used the link Nearest Neighbor Calculator for Nucleic Acids (konan-fiber.jp) for these calculations[19–22].

### 4.12. Structure preparation and molecular dynamics simulation

SpCas9-sgRNA-DNA ternary complex structure was taken from PDB (pdb id: 5Y36) and any missing atoms were added using CHARMM-GUI server [53]. We have replaced the original guideRNA and target DNA sequence of the pdb structure with our own guideRNA and target DNA sequence using w3DNA 2.0 webserver [54]. This system is denoted as Wild type. Then several mismatch mutations (between RNA-DNA hybrid) in the target DNA strands of wild type were introduced using the same server resulting in 23 independent systems with wild type system and 22 different mismatch DNA sequences (**Figure1e**).Molecular dynamics simulation was conducted for total 23 systems. GROMACS 2022 [55] engine was used for the all the simulations. AMBER99-IILDN [56] force field was used in this study and the temperature was set to 300 K. All the systems were placed in a dodecahedron box with box edge of 12Å and 8Å distance was set between protein atoms and box edge. All the systems were solvated using TIP3 water models and the solvated system were neutralized by adding sufficient numbers of Na+ and Cl-counter ions. All the systems were energy minimized using steepest descent algorithm with 50000 steps. The minimzation step is continued until the maximum force on the atoms is less than 1000 kJ/mol/nm. Then we performed 100 ps of NVT equilibration step in order to equilibrate the system at 300 K temperature using the Langevin thermostat [57]and subsequently 100 ps of NPT equilibration in order to equilibrate it at 1 atm pressure using Berendsen barostat[58]. During the NVT and NPT equilibration all the heavy atom were restrained with a positional constraint of 1000 kcal/mol Å-2. Production run for each system was conducted for 50 ns in the NPT ensemble with no restraint resulting in total 1000 ns of simulation. To handle long-range Coulomb interactions, the particle mesh Ewald summation method (PME) [59] was employed by setting the mesh spacing to 1.0 Å. All the analysis were done using inbuild gromacs command line tool and plotting was done using Python matplotlib module.

## 5. Acknowledgments

The authors would like to thank Dr.Sejuti Sinha Roy, Dr. Dhananjay Bhattacharya, Sayandeep Gupta, Debaleena Bhowmik, and Arina Yasmin for the discussion on molecular biology concept, nearest-neighbor model approach, and data sharing by mega. NZ. The authors would like to thank the **Department of Biotechnology, India** (**DBT)** for funding the project titled **”Exploitation Of Paired Nickases To Enhance Targeted Knock-In: A Modification Of Crispr Cas System” and project sanction number BT/PR 26301/GET/119/258/2017.** RC would also like to acknowledge **DST-INSPIRE** for providing him fellowship. SR would like to acknowledge **ICMR** for granting him Emeritus position.

## 6. Conflict of Interest

The authors declare there is no conflict of interest.

## 7. Author contribution role

DD & DG-designed the experiments

DD, RC, OB, RS& RG– Worked on the experiments

DD, RC, SM, DC, AG, & RS – In-silico experiments

VR, DC, AG, KTA, SR & DG-Critical Review of the paper

## 8. Data availability

Authors agree to provide materials, used in our study, promptly available to others upon request.

## Notes

### Competing Interest Statement

The authors have declared no competing interest.

## Bibliography

1. Gupta, D., et al., CRISPR-Cas9 system: A new-fangled dawn in gene editing. Life sciences, 2019. 232: p. 116636.

2. Wu, X., A.J. Kriz, and P.A. Sharp, Target specificity of the CRISPR-Cas9 system. Quantitative biology, 2014. 2: p. 59–70.

3. Gupta, R., et al., Modification of Cas9, gRNA and PAM: key to further regulate genome editing and its applications. Progress in Molecular Biology and Translational Science, 2021. 178: p. 85–98.

4. Nishimasu, H., et al., Crystal structure of Cas9 in complex with guide RNA and target DNA. Cell, 2014. 156(5): p. 935–949.

5. Wong, N., W. Liu, and X. Wang, WU-CRISPR: characteristics of functional guide RNAs for the CRISPR/Cas9 system. Genome biology, 2015. 16(1): p. 1–8.

6. Cong, L., et al., Multiplex genome engineering using CRISPR/Cas systems. Science, 2013. 339(6121): p. 819–823.

7. Jiang, W., et al., RNA-guided editing of bacterial genomes using CRISPR-Cas systems. Nature biotechnology, 2013. 31(3): p. 233–239.

8. Cho, S.W., et al., Analysis of off-target effects of CRISPR/Cas-derived RNA-guided endonucleases and nickases. Genome research, 2014. 24(1): p. 132–141.

9. Hsu, P.D., et al., DNA targeting specificity of RNA-guided Cas9 nucleases. Nature biotechnology, 2013. 31(9): p. 827–832.

10. Bravo, J.P.K., et al., Structural basis for mismatch surveillance by CRISPR-Cas9. Nature, 2022. 603(7900): p. 343–347.

11. Pacesa, M., et al., Structural basis for Cas9 off-target activity. Cell, 2022. 185(22): p. 4067–4081.e21.

12. Wang, T., et al., Genetic screens in human cells using the CRISPR-Cas9 system. Science, 2014. 343(6166): p. 80–84.

13. Huai, C., et al., Structural insights into DNA cleavage activation of CRISPR-Cas9 system. Nat Commun, 2017. 8(1): p. 1375.

14. Jinek, M., et al., A programmable dual-RNA-guided DNA endonuclease in adaptive bacterial immunity. Science, 2012. 337(6096): p. 816–21.

15. Gasiunas, G., et al., Cas9-crRNA ribonucleoprotein complex mediates specific DNA cleavage for adaptive immunity in bacteria. Proc Natl Acad Sci U S A, 2012. 109(39): p. E2579–86.

16. Chen, H., J. Choi, and S. Bailey, Cut site selection by the two nuclease domains of the Cas9 RNA-guided endonuclease. J Biol Chem, 2014. 289(19): p. 13284–94.

17. Fu, Y., et al., Improving CRISPR-Cas nuclease specificity using truncated guide RNAs. Nature biotechnology, 2014. 32(3): p. 279–284.

18. Fu, B.X., et al., Distinct patterns of Cas9 mismatch tolerance in vitro and in vivo. Nucleic Acids Research, 2016. 44(11): p. 5365–5377.

19. SantaLucia, J., H.T. Allawi, and P.A. Seneviratne, Improved nearest-neighbor parameters for predicting DNA duplex stability. Biochemistry, 1996. 35(11): p. 3555–3562.

20. Sugimoto, N., et al., Thermodynamic parameters to predict stability of RNA/DNA hybrid duplexes. Biochemistry, 1995. 34(35): p. 11211–11216.

21. Freier, S.M., et al., Improved free-energy parameters for predictions of RNA duplex stability. Proceedings of the National Academy of Sciences, 1986. 83(24): p. 9373–9377.

22. Banerjee, D., et al., Improved nearest-neighbor parameters for the stability of RNA/DNA hybrids under a physiological condition. Nucleic Acids Research, 2020. 48(21): p. 12042–12054.

23. Yang, H., et al., One-step generation of mice carrying reporter and conditional alleles by CRISPR/Cas-mediated genome engineering. Cell, 2013. 154(6): p. 1370–1379.

24. Long, C., et al., Postnatal genome editing partially restores dystrophin expression in a mouse model of muscular dystrophy. Science, 2016. 351(6271): p. 400-403.

25. Kim, H.J., et al., Targeted genome editing in human cells with zinc finger nucleases constructed via modular assembly. Genome research, 2009. 19(7): p. 1279–1288.

26. Brunet, E., et al., Chromosomal translocations induced at specified loci in human stem cells. Proceedings of the National Academy of Sciences, 2009. 106(26): p. 10620–10625.

27. Lee, H.J., et al., Targeted chromosomal duplications and inversions in the human genome using zinc finger nucleases. Genome research, 2012. 22(3): p. 539–548.

28. Lee, H.J., E. Kim, and J.-S. Kim, Targeted chromosomal deletions in human cells using zinc finger nucleases. Genome research, 2010. 20(1): p. 81–89.

29. Kim, Y., et al., A library of TAL effector nucleases spanning the human genome. Nature biotechnology, 2013. 31(3): p. 251–258.

30. Ivanov, I.E., et al., Cas9 interrogates DNA in discrete steps modulated by mismatches and supercoiling. Proceedings of the National Academy of Sciences, 2020. 117(11): p. 5853–5860.

31. Anderson, E.M., et al., Systematic analysis of CRISPR–Cas9 mismatch tolerance reveals low levels of off-target activity. Journal of biotechnology, 2015. 211: p. 56–65.

32. Hewes, A.M., et al., gRNA sequence heterology tolerance catalyzed by CRISPR/Cas in an in vitro homology-directed repair reaction. Molecular Therapy-Nucleic Acids, 2020. 20: p. 568–579.

33. Chuai, G., et al., DeepCRISPR: optimized CRISPR guide RNA design by deep learning. Genome Biol, 2018. 19(1): p. 80.

34. Haeussler, M., et al., Evaluation of off-target and on-target scoring algorithms and integration into the guide RNA selection tool CRISPOR. Genome Biol, 2016. 17(1): p. 148.

35. Listgarten, J., et al., Prediction of off-target activities for the end-to-end design of CRISPR guide RNAs. Nat Biomed Eng, 2018. 2(1): p. 38–47.

36. Singh, R., et al., Cas9-chromatin binding information enables more accurate CRISPR off-target prediction. Nucleic Acids Res, 2015. 43(18): p. e118.

37. Stemmer, M., et al., Correction: CCTop: An Intuitive, Flexible and Reliable CRISPR/Cas9 Target Prediction Tool. PLoS One, 2017. 12(4): p. e0176619.

38. Yan, J., et al., Benchmarking and integrating genome-wide CRISPR off-target detection and prediction. Nucleic Acids Res, 2020. 48(20): p. 11370–11379.

39. Zhang, D., et al., Unified energetics analysis unravels SpCas9 cleavage activity for optimal gRNA design. Proc Natl Acad Sci U S A, 2019. 116(18): p. 8693–8698.

40. Zhao, C., et al., CRISPR-offinder: a CRISPR guide RNA design and off-target searching tool for user-defined protospacer adjacent motif. Int J Biol Sci, 2017. 13(12): p. 1470–1478.

41. Dagdas, Y.S., et al., A conformational checkpoint between DNA binding and cleavage by CRISPR-Cas9. Science advances, 2017. 3(8): p. eaao0027.

42. Zhang, X.-H., et al., Off-target effects in CRISPR/Cas9-mediated genome engineering. Molecular Therapy-Nucleic Acids, 2015. 4.

43. Yan, J., et al., Benchmarking and integrating genome-wide CRISPR off-target detection and prediction. Nucleic acids research, 2020. 48(20): p. 11370–11379.

44. Stemmer, M., et al., CCTop: an intuitive, flexible and reliable CRISPR/Cas9 target prediction tool. PloS one, 2015. 10(4): p. e0124633.

45. Singh, R., et al., Cas9-chromatin binding information enables more accurate CRISPR off-target prediction. Nucleic acids research, 2015. 43(18): p. e118–e118.

46. Haeussler, M., et al., Evaluation of off-target and on-target scoring algorithms and integration into the guide RNA selection tool CRISPOR. Genome biology, 2016. 17: p. 1–12.

47. Zhao, C., et al., CRISPR-offinder: a CRISPR guide RNA design and off-target searching tool for user-defined protospacer adjacent motif. International journal of biological sciences, 2017. 13(12): p. 1470.

48. Alkan, F., et al., CRISPR-Cas9 off-targeting assessment with nucleic acid duplex energy parameters. Genome biology, 2018. 19(1): p. 1–13.

49. Listgarten, J., et al., Prediction of off-target activities for the end-to-end design of CRISPR guide RNAs. Nature biomedical engineering, 2018. 2(1): p. 38–47.

50. Chuai, G., et al., DeepCRISPR: optimized CRISPR guide RNA design by deep learning. Genome biology, 2018. 19: p. 1–18.

51. Zhang, D., et al., Unified energetics analysis unravels SpCas9 cleavage activity for optimal gRNA design. Proceedings of the National Academy of Sciences, 2019. 116(18): p. 8693–8698.

52. Ricci, C.G., et al., Deciphering off-target effects in CRISPR-Cas9 through accelerated molecular dynamics. ACS central science, 2019. 5(4): p. 651–662.

53. Jo, S., et al., CHARMM-GUI 10 years for biomolecular modeling and simulation. J Comput Chem, 2017. 38(15): p. 1114–1124.

54. Li, S., W.K. Olson, and X.J. Lu, Web 3DNA 2.0 for the analysis, visualization, and modeling of 3D nucleic acid structures. Nucleic Acids Res, 2019. 47(W1): p. W26–w34.

55. Abraham, M.J., et al., GROMACS: High performance molecular simulations through multi-level parallelism from laptops to supercomputers. SoftwareX, 2015. 1: p. 19–25.

56. Lindorff-Larsen, K., et al., Improved side-chain torsion potentials for the Amber ff99SB protein force field. Proteins, 2010. 78(8): p. 1950–8.

57. Farago, O., Langevin thermostat for robust configurational and kinetic sampling. Physica A: Statistical Mechanics and Its Applications, 2019. 534: p. 122210.

58. Berendsen, H.J., et al., Molecular dynamics with coupling to an external bath. The Journal of chemical physics, 1984. 81(8): p. 3684–3690.

59. Darden, T., D. York, and L. Pedersen, Particle mesh Ewald: An N⋅ log (N) method for Ewald sums in large systems. The Journal of chemical physics, 1993. 98(12): p. 10089–10092.

