## supplementary file 1 for "A mechanistic study on the tolerance of PAM distal end mismatch by SpCas9"

**SUPPLEMENTARY TABLE**

**Table A-single mismatches**

| **NAME** | **SEQUENCE (5’ > 3’)** | **ΔG^o^ (kcal mol−1)** |
| --- | --- | --- |
| **WT** | **GCTTCTACCCCAATGACTTG** | **-17.8** |
| **20MM** | **ACTTCTACCCCAATGACTTG** | **-15.8** |
| **19MM** | **GTTTCTACCCCAATGACTTG** | **-16.8** |
| **18MM** | **GCCTCTACCCCAATGACTTG** | **-18** |
| **17MM** | **GCTCCTACCCCAATGACTTG** | **-18** |
| **16MM** | **GCTTTTACCCCAATGACTTG** | **-15.8** |
| **15MM** | **GCTTCCACCCCAATGACTTG** | **-18.1** |
| **14MM** | **GCTTCTGCCCCAATGACTTG** | **-17.2** |
| **13MM** | **GCTTCTATCCCAATGACTTG** | **-16.1** |
| **12MM** | **GCTTCTACTCCAATGACTTG** | **-.16.1** |

**Table *B* -Bi-Sequential Mismatch**

| **NAME** | **SEQUENCE (5’ > 3’)** | **ΔG^o^ (kcal mol−1)** |
| --- | --- | --- |
| **WT** | **GCTTCTACCCCAATGACTTG** | **-17.8** |
| **2019MM** | **ATTTCTACCCCAATGACTTG** | **-15.4** |
| **1918MM** | **GTCTCTACCCCAATGACTTG** | **-17** |
| **1817MM** | **GCCCCTACCCCAATGACTTG** | **-17.9** |
| **1716MM** | **GCTCTTACCCCAATGACTTG** | **-16** |
| **1615MM** | **GCTTTCACCCCAATGACTTG** | **-16** |
| **1514MM** | **GCTTCCGCCCCAATGACTTG** | **-17.1** |
| **1413MM** | **GCTTCTGTCCCAATGACTTG** | **-15.5** |
| **1312MM** | **GCTTCTATTCCAATGACTTG** | **-14.4** |

**Table *C*- Sequential Mismatches and triple mismatches**

| **NAME** | **SEQUENCE (5’ > 3’)** | **ΔG^o^ (kcal mol−1)** |
| --- | --- | --- |
| **WT** | **GCTTCTACCCCAATGACTTG** | **-17.8** |
| **2018MM** | **ATCTCTACCCCAATGACTTG** | **-15.6** |
| **2017MM** | **ATCCCTACCCCAATGACTTG** | **-14.2** |
| **2016MM** | **ATCCTTACCCCAATGACTTG** | **-13.8** |
| **2015MM** | **ATCCTCACCCCAATGACTTG** | **-13.3** |
| **1917MM** | **GTCCCTACCCCAATGACTTG** | **-16.2** |
| **1816MM** | **GCCCTTACCCCAATGACTTG** | **-16.2** |
| **1715MM** | **GCTCTCACCCCAATGACTTG** | **-16.2** |

**Table *D*- Staggered Mismatches**

| **NAME** | **SEQUENCE (5’ > 3’)** | **ΔG^o^ (kcal mol−1)** |
| --- | --- | --- |
| **WT** | **GCTTCTACCCCAATGACTTG** | **-17.8** |
| **20/18MM** | **ACCTCTACCCCAATGACTTG** | **-16** |
| **20/18/16MM** | **ACCTTTACCCCAATGACTTG** | **-14** |
| **20/18/16/14MM** | **ACCTTTGCCCCAATGACTTG** | **-13.4** |
| **20/18/16/14/12MM** | **ACCTTTGCTCCAATGACTTG** | **-11.7** |
| **20/16MM** | **ACTTTTACCCCAATGACTTG** | **-13.8** |
| **20/14MM** | **ACTTCTGCCCCAATGACTTG** | **-15.2** |
| **18/16MM** | **GCCTTTACCCCAATGACTTG** | **-16** |
| **18/14MM** | **GCCTCTGCCCCAATGACTTG** | **-17.4** |
| **16/14MM** | **GCTTTTGCCCCAATGACTTG** | **-15.2** |
| **20/1817MM** | **ACCCCTACCCCAATGACTTG** | **-15.9** |
| **20/1716MM** | **ACTCTTACCCCAATGACTTG** | **-14** |
| **20/1615MM** | **ACTTTCACCCCAATGACTTG** | **-14** |
| **20/18/1615MM** | **ACCTTCACCCCAATGACTTG** | **-14.2** |
| **18/1615MM** | **GCCTTCACCCCAATGACTTG** | **-16.2** |
| **18/1514MM** | **GCCTCCGCCCCAATGACTTG** | **-17.3** |
