## supplementary file 2 for "A mechanistic study on the tolerance of PAM distal end mismatch by SpCas9"

1. **Negative**
2. **GCGTACGACATCGCAGACTG- TS5 RNA 20**
3. **CGTACGACATCGCAGACTG- TS5 RNA 19**
4. **GTACGACATCGCAGACTG- TS5 RNA 18**
5. **TACGACATCGCAGACTG- TS5 RNA 17**
6. **ACGACATCGCAGACTG- TS5 RNA 16**
7. **CGACATCGCAGACTG- TS5 RNA 15**

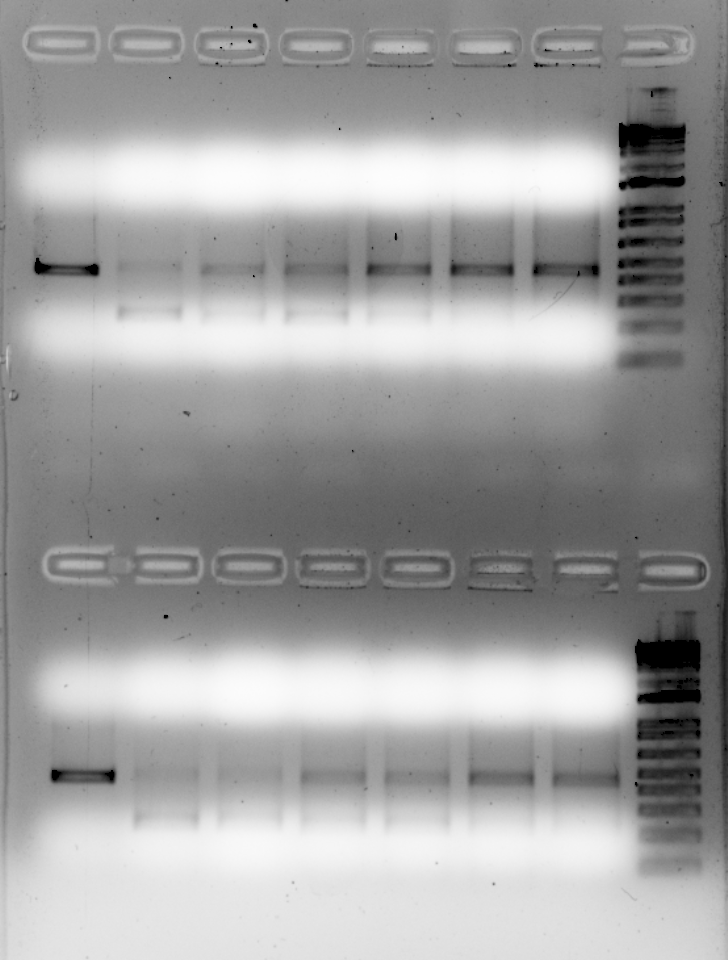

**1	2	3	4	5	6	7**

1. **Negative**
2. **GCTTACAACATTGTGAACGA- TS4 RNA 20**
3. **CTTACAACATTGTGAACGA- TS4 RNA 19**
4. **TTACAACATTGTGAACGA- TS4 RNA 18**
5. **TACAACATTGTGAACGA- TS4 RNA 17**
6. **ACAACATTGTGAACGA- TS4 RNA 16**
7. **CAACATTGTGAACGA- TS4 RNA 15**

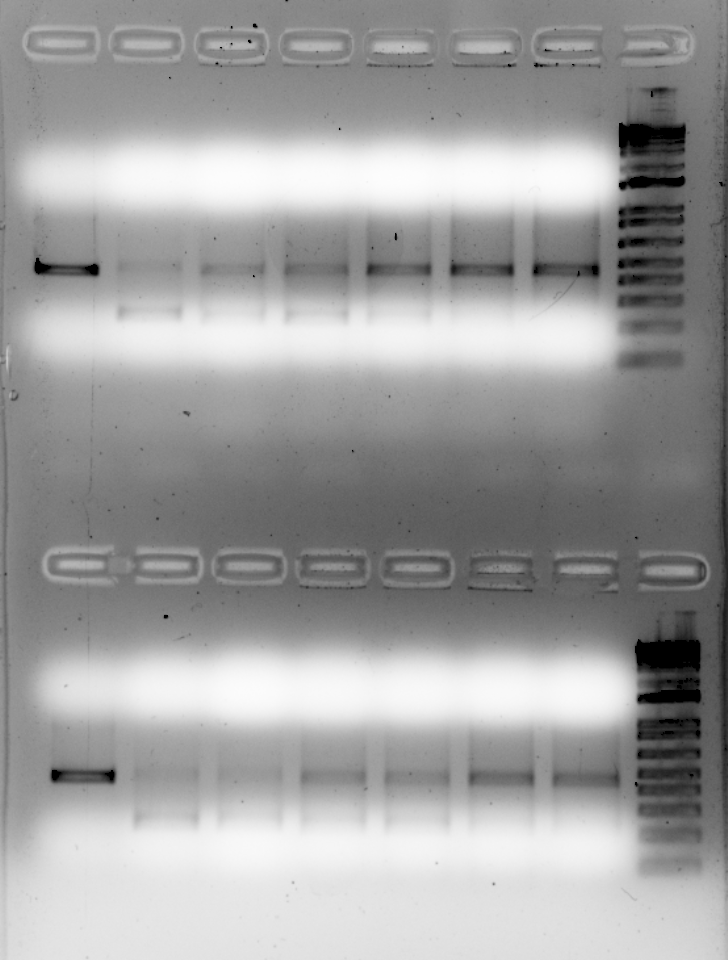

**1	2	3	4	5	6	7**

Table 1- Complementary binding energy due to RNA drop

| **NAME** | **SEQUENCE 5’-3’** | **ΔG^o^ (kcal mol−1)** |
| --- | --- | --- |
| **TS4 RNA 20** | **GCUUACAACAUUGUGAACGA** | **-18.2** |
| **TS4 RNA 19** | **CUUACAACAUUGUGAACGA** | **-16.2** |
| **TS4 RNA 18** | **UUACAACAUUGUGAACGA** | **-15.2** |
| **TS4 RNA 17** | **UACAACAUUGUGAACGA** | **-15.4** |
| **TS4 RNA 16** | **ACAACAUUGUGAACGA** | **-14.9** |
| **TS4 RNA 15** | **CAACAUUGUGAACGA** | **-14** |
| **TS5 RNA 20** | **GCGUACGACAUCGCAGACUG** | **-23.1** |
| **TS5 RNA 19** | **CGUACGACAUCGCAGACUG** | **-21.1** |
| **TS5 RNA 18** | **GUACGACAUCGCAGACUG** | **-19.7** |
| **TS5 RNA 17** | **UACGACAUCGCAGACUG** | **-18.3** |
| **TS5 RNA 16** | **ACGACAUCGCAGACUG** | **-17.8** |
| **TS5 RNA 15** | **CGACAUCGCAGACUG** | **-16.3** |

*DNA digestion with truncated gRNA probe TS4 and TS5*

1. **Negative**
2. **GCGTACGACATCGCAGACTG- TS5 RNA 20**
3. **CGTACGACATCGCAGACTG- TS5 RNA 19**
4. **GTACGACATCGCAGACTG- TS5 RNA 18**
5. **TACGACATCGCAGACTG- TS5 RNA 17**
6. **ACGACATCGCAGACTG- TS5 RNA 16**
7. **CGACATCGCAGACTG- TS5 RNA 15**
